# Discovery of widespread transcription initiation at microsatellites predictable by sequence-based deep neural network

**DOI:** 10.1101/2020.07.10.195636

**Authors:** Mathys Grapotte, Manu Saraswat, Chloé Bessière, Christophe Menichelli, Jordan A. Ramilowski, Jessica Severin, Yoshihide Hayashizaki, Masayoshi Itoh, Michihira Tagami, Mitsuyoshi Murata, Miki Kojima-Ishiyama, Shohei Noma, Shuhei Noguchi, Takeya Kasukawa, Akira Hasegawa, Harukazu Suzuki, Hiromi Nishiyori-Sueki, Martin C. Frith, FANTOM consortium, Clément Chatelain, Piero Carninci, Michiel J.L. de Hoon, Wyeth W. Wasserman, Laurent Bréhélin, Charles-Henri Lecellier

## Abstract

Using the Cap Analysis of Gene Expression (CAGE) technology, the FANTOM5 consortium provided one of the most comprehensive maps of Transcription Start Sites (TSSs) in several species. Strikingly, ~ 72% of them could not be assigned to a specific gene and initiate at unconventional regions, outside promoters or enhancers. Here, we probed these unassigned TSSs and showed that, in all species studied, a significant fraction of CAGE peaks initiate at microsatellites, also called short tandem repeats (STRs). To confirm this transcription, we developed Cap Trap RNA-seq, a technology which combines cap trapping and long reads MinION sequencing. We trained sequence-based deep learning models able to predict CAGE signal at STRs with high accuracy. These models unveiled the importance of STR surrounding sequences not only to distinguish STR classes, as defined by the repeated DNA motif, one from each other, but also to predict their transcription. Excitingly, our models predicted that genetic variants linked to human diseases affect STR-associated transcription and correspond precisely to the key positions identified by our models to predict transcription. Together, our results extend the repertoire of non-coding transcription associated with DNA tandem repeats and complexify STR polymorphism.

## Introduction

RNA polymerase II (RNAPII) transcribes many loci outside annotated protein-coding gene promoters ^1, 2^ to generate a diversity of RNAs, including for instance enhancer RNAs ^3^ and long noncoding RNAs ^4^. In fact, > 70% of all nucleotides are thought to be transcribed at some point^1, 5, 6^. Using the Cap Analysis of Gene Expression (CAGE) technology ^7, 8^, the FANTOM5 consortium provided one of the most comprehensive maps of TSSs in several species ^2^. Integrating multiple collections of transcript models with FANTOM CAGE datasets, Hon *et al*. built a new annotation of the human genome (FANTOM CAGE Associated Transcriptome, FANTOM CAT), with an atlas of 27,919 human lncRNAs, among them 19,175 potentially functional RNAs ^4^. Despite this annotation, many CAGE peaks remain unassigned to a specific gene and/or initiate at unconventional regions, outside promoters or enhancers, providing an unprecedented mean to further characterize non-coding transcription within the genome ‘dark matter’ ^9^ and to decode part of the transcriptional noise.

Non-coding transcription is indeed far from being fully understood ^10^ and some authors suggest that many of these transcripts, often faintly expressed, can simply be ‘noise’ or ‘junk’ ^11, 12^. On the other hand, many non annotated RNAPII transcribed regions correspond to open chromatin ^1^ and *cis*-regulatory modules bound by transcription factors (TFs) ^13^. Besides, genome-wide association studies showed that trait-associated loci, including those linked to human diseases, can be found outside canonical gene regions^14–16^. Together, these findings suggest that the non-coding regions of the human genome harbor a plethora of potentially transcribed functional elements, which can drastically impact genome regulations and functions ^9, 16^.

The human genome is scattered with repetitive sequences, and a large portion of non-coding RNAs derive from repetitive elements ^17, 18^, in particular DNA tandem repeats, such as satellite DNAs ^19^ and minisatellites ^20^. Microsatellites, also called Short tandem repeats (STRs), constitute a third class of DNA tandem repeats. They correspond to repeated DNA motifs of 2 to 6 bp and constitute one of the most polymorphic and abundant repetitive elements ^21^. Classes of STRs can be defined based on the repeated DNA motif (e.g. (*AC*)_*n*_ will correspond to all STRs with repeats of the dinucleotide AC). STR polymorphism, which corresponds to variation in number of repeated DNA motif (i.e. STR length), is presumably due to their susceptibility to slippage events during DNA replication. STRs have been shown to widely impact gene expression and to contribute to expression variation ^22–24^. Some constitute genuine expression Quantitative Trait Loci (eQTLs) ^22, 23^, called eSTRs ^22^. At the molecular level, STR can for instance affect expression by inducing inhibitory DNA structures ^25^ and/or by modulating the binding of transcription factors ^26, 27^.

Provided the abundance of STRs on the one hand and the widespread transcription of the genome, including at repeated elements, on the other hand, we hypothesized that transcription initiation also occurs at STRs. To test this hypothesis, we probed CAGE data collected by the FANTOM5 consortium ^2^ using the STRs catalog built by Willems *et al*. ^28^. We specifically showed that a significant portion of CAGE peaks (~ 8.6%) initiate at STRs. This transcription, which was confirmed by Cap Trap RNA-seq (CTR-seq), a technology which combines cap trapping and long read MinIoN sequencing, not only extends the repertoire of non-coding transcription but also STR polymorphism, with STRs having the same length but different transcription rate and, conversely, STRs with different lengths having similar transcription rate. We further learned a sequencebased Convolutional Neural Networks (CNNs) able to predict this transcription with high accuracy (correlation between observed and predicted CAGE signal > 0.68 for 14 STR classes with > 5,000 elements). These models unveil the importance of STR flanking sequences in distinguishing STR classes, one from each other, and also in predicting their transcription. We finally showed that many genetic variants linked to human diseases, including those located, not only within, but also around STRs, can affect this STR-associated transcription, thereby advancing our capacity to interpret several nucleotide variants.

## Results

### CAGE peaks are detected at STRs

We first intersected the coordinates of 1,048,124 CAGE peak summits ^2^ with that of 1,620,030 STRs called by HipSTR ^28^. We found that 89,948 CAGE peaks (~ 8.6%) initiate at 84,555 STRs (Figure 1A and Supplementary Figure S1). Among these CAGE peaks, 10,727 correspond to TSSs of FANTOM CAT transcripts^4^ and 8,823 to enhancer boundaries ^3^ (Supplementary Table S1). Note that the FANTOM CAT annotation was shown to be more accurate in 5’ end transcript definitions compared to other catalogs, because transcript models combine various independent sources (GENCODE release 19, Human BodyMap 2.0, mi-Transcriptome, ENCODE and an RNA-seq assembly from 70 FANTOM5 samples) and FANTOM CAT TSSs were validated with Roadmap Epigenome DHS and RAMPAGE data sets ^4^. This transcription does not correspond to random noise because the fraction of STRs harbouring a CAGE peak within each class differs depending on the STR class, without any link with their abundance (Figures 1B and 1C). Some STR classes with low abundance are indeed more often associated with a CAGE peak than more abundant STRs (Figures 1B and 1C, compare for instance (*CTTTTT*)_*n*_ or (*AAAAG*)_*n*_ vs. (*AT*)_*n*_ or (*ATTT*)_*n*_). Likewise, the number of STRs associated with CAGE peaks cannot merely be explained by their length, as several STR classes have similar length distribution but very different fractions of CAGE-associated loci (compare for instance (*AT*)_*n*_ and (*GT*)_*n*_ in Figure 1C and Supplementary Figure S2).

**Figure 1.**
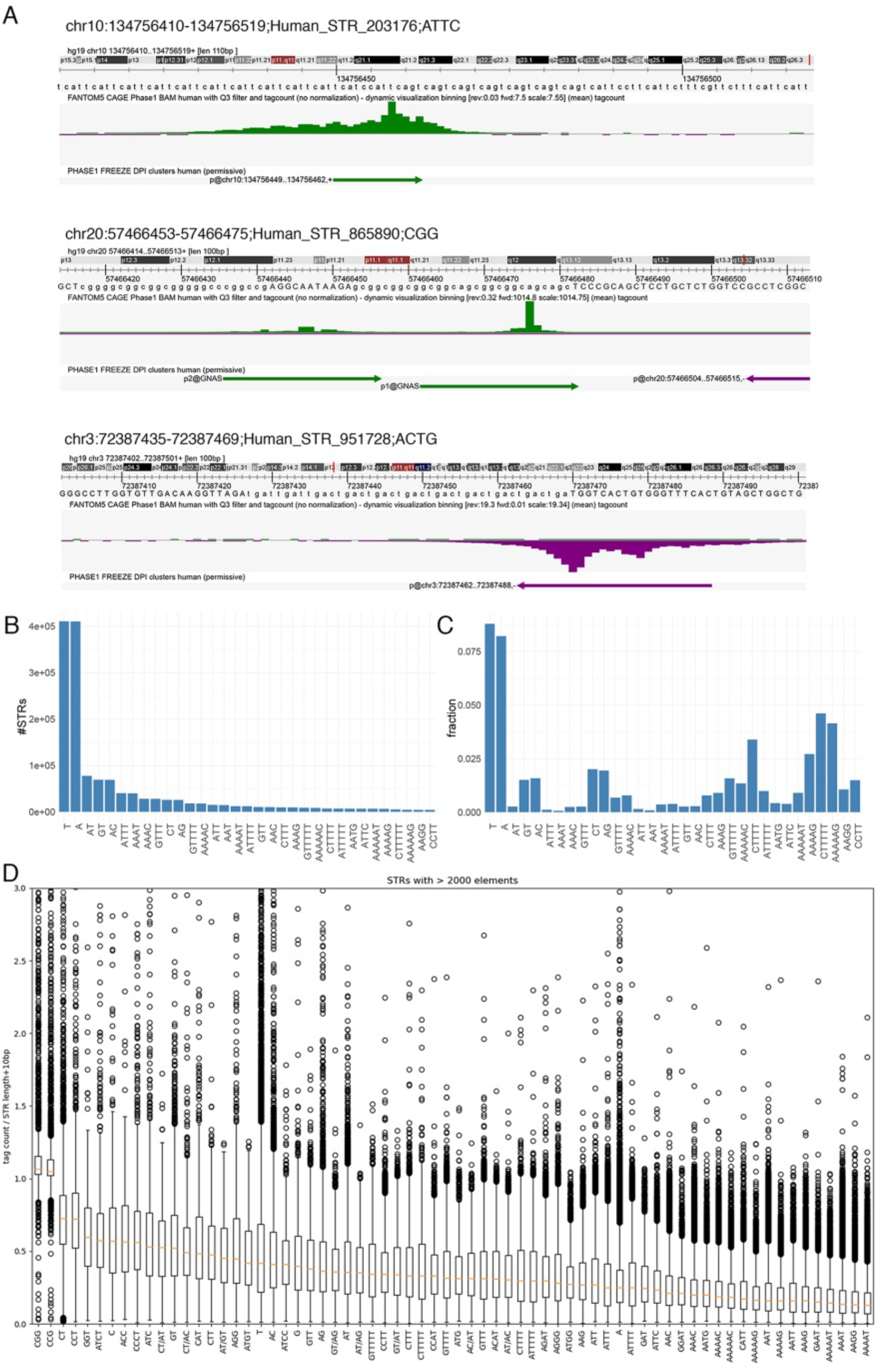
CAGE peaks are detected at STRs. **A.** Three examples of STRs associated with a CAGE peak. The Zenbu browser^62^ was used. top track, hg19 genome sequence; middle track, CAGE tag count as mean across 988 libraries (BAM files with Q3 filter were used); bottom track, CAGE peaks as called in ^2^. **B.** Number of STRs per STR class. For sake of clarity, only STR classes with > 2,000 loci are shown. **C.** Fraction of STRs associated with a CAGE peak in all STR classes considered in B. **D.** CAGE signal at STR classes with > 2,000 loci. CAGE signal was computed as the mean raw tag count of each STR (tag count in STR ± 5bp) across all 988 FANTOM5 libraries. This tag count was further normalized by the length of the window used to compute the signal (i.e. STR length + 10bp).

We next computed the tag count sum along each STR ± 5bp, and averaged the signal across 988 FANTOM5 libraries. We noticed the existence of very low (tagcount = 1) CAGE tag counts all along STRs, which artificially increase the signal (see examples in Figure 1A, Spearman correlation coefficient between sum CAGE tag count along STR and STR length ~ 0.26). To remove any dependence between STR length and CAGE signal, the mean tag count was therefore normalized by the length of the window used to compute the signal (i.e. STR length + 10bp). Looking directly at this CAGE signal (not CAGE peaks) along the genome, we observed that some STR classes are more transcribed than others (Figure 1D, compare (*CGG*)_*n*_ or (*CCG*)_*n*_ vs. (*AAGG*)_*n*_ or (*AAAAT*)_*n*_). While STRs are mostly intragenic (1,195,065 out of 1,620,030), there is no drastic difference in term of transcription between intra- and intergenic STRs (Supplementary Figure S3). Looking at each STR class separately, we confirmed that our CAGE signal computation is not sensitive to the STR length (Supplementary Figure S4). Supplementary Figure S4 also shows that STRs with different lengths can be associated with the same CAGE signal while, conversely, that two STRs with different CAGE signals can have the same length. Thus, considering transcription, STR polymorphism appears to not only rely on the number of repeated elements. Transcription therefore appears to complexify STR polymorphism.

### CAGE tags correspond to genuine transcriptional products

CAGE read detection at STRs faces two problems. First, CAGE tags can capture not only TSSs but also the 5’ ends of post-transcriptionally processed RNAs ^29^. To clarify that point, we used a strategy described by de Rie *et al*. ^30^, which compares CAGE tags obtained by Illumina (ENCODE) vs. Heliscope (FANTOM) technologies. Briefly, the 7-methylguanosine cap at the 5’ end of CAGE tags produced by RNA polymerase II can be recognized as a guanine nucleotide during reverse transcription. This artificially introduces mismatched Gs at Illumina tag 5’ end, which is not detected with Heliscope sequencing, which skips the first nucleotide ^30^. We then evaluated the existence of this G bias in CAGE tags corresponding to peaks detected at STRs, peaks assigned to genes (for positive control), and peaks intersecting the 3’ end of precursor microRNAs (pre-miRNAs for negative control) (Figure 2). While most CAGE tag 5’ ends perfectly match the sequences of pre-miRNA 3’end in all cell types tested, as previously reported ^30^, a G bias was clearly observed when considering assigned CAGEs and CAGEs detected at STRs, confirming that the vast majority of STR-associated CAGE tags are truly capped. We also confirmed that STRs located within RNAPII binding sites exhibit a stronger CAGE signal than STRs not associated with RNAPII binding events (Supplementary Figure S5).

**Figure 2.**
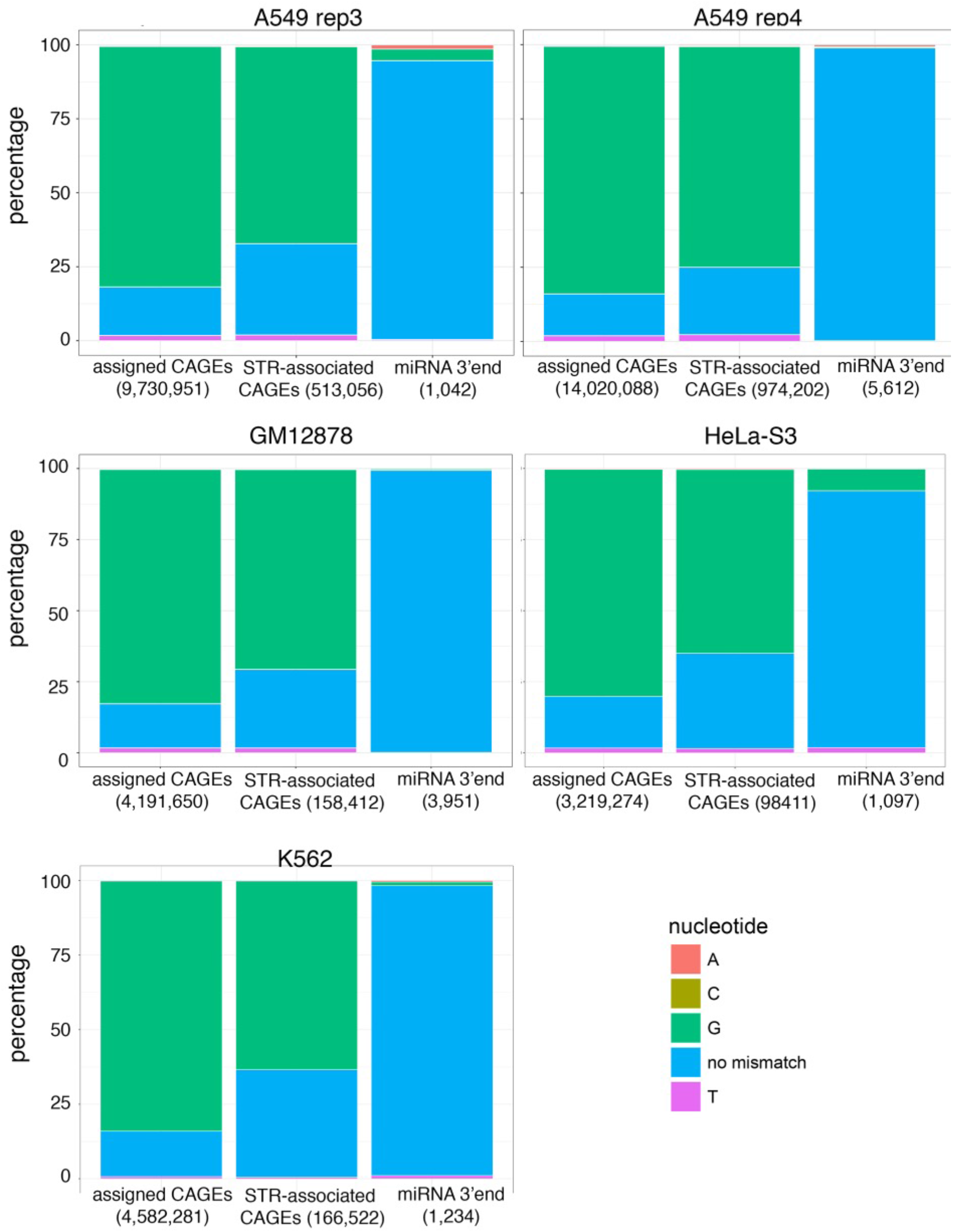
CAGE tags initiating at STRs are truly 5’-capped. G bias in ENCODE CAGE tags (bam files from nuclear fraction, polyA+) was assessed at FANTOM5 CAGE peaks assigned to genes (positive control) and CAGE peaks initiating at STRs. G bias at pre-microRNA 3’ ends was also assessed as negative control. Five libraries were analyzed corresponding to A549 (replicates 3 and 4), GM12878, HeLa-S3 and K562 cells. The number of intersecting tags in each case is indicated in bracket.

Second, because of their repetitive nature, mapping CAGE reads to STRs is problematic and may yield ambiguous results. To circumvent this issue, we used CTR-seq, which combines cap trapping and long read MinION sequencing. With this technology, median read length is > 500 bp, thereby greatly limiting the chance of erroneous mapping. Two libraries were generated in A549 cells, including or not polyA tailing. This polyA tailing step before reverse transcription allows the detection of polyA-minus non-coding RNAs. Long reads initiating at STRs were readily detected in both libraries (Figures 3A and 3B). As expected provided the depth of MinION sequencing in only one cell line, the number of STRs associated with long reads is lower than that obtained with CAGE sequencing collected in 988 libraries (n = 5,472 and 7,812, respectively with and without polyA tailing with 2,291 STRs associated with long reads in both libraries). Among these 2,291 STRs, 904 (39%) are also associated with a CAGE peak. Thus, compared to the reproducibility of MinION sequencing in both libraries (only 2,291 STRs in common out of 5,472 (42%) or 7,812 (29%)), CAGE and CTR-seq sequencing results are overall in agreement. In fact, STR classes associated with CAGE peaks correspond to that associated with CTR-seq reads (Figures 3A and B compared to Figure 1C). The Spearman correlation *ρ* between the fractions of STRs associated with CAGE and MinION reads with and without polyA tailing equals 0.88 and 0.89 respectively. Besides, 301 out of 904 STRs associated with both CAGE peak and CTR-seq long read correspond to TSSs of FANTOM CAT transcripts and 54 to enhancer boundaries. Overall, CTR-seq confirms CAGE data and the existence of transcription initiating at STRs. The similarity of the results obtained with and without the polyA tailing step also indicates that RNAs initiating at STRs are mostly polyadenylated.

**Figure 3.**
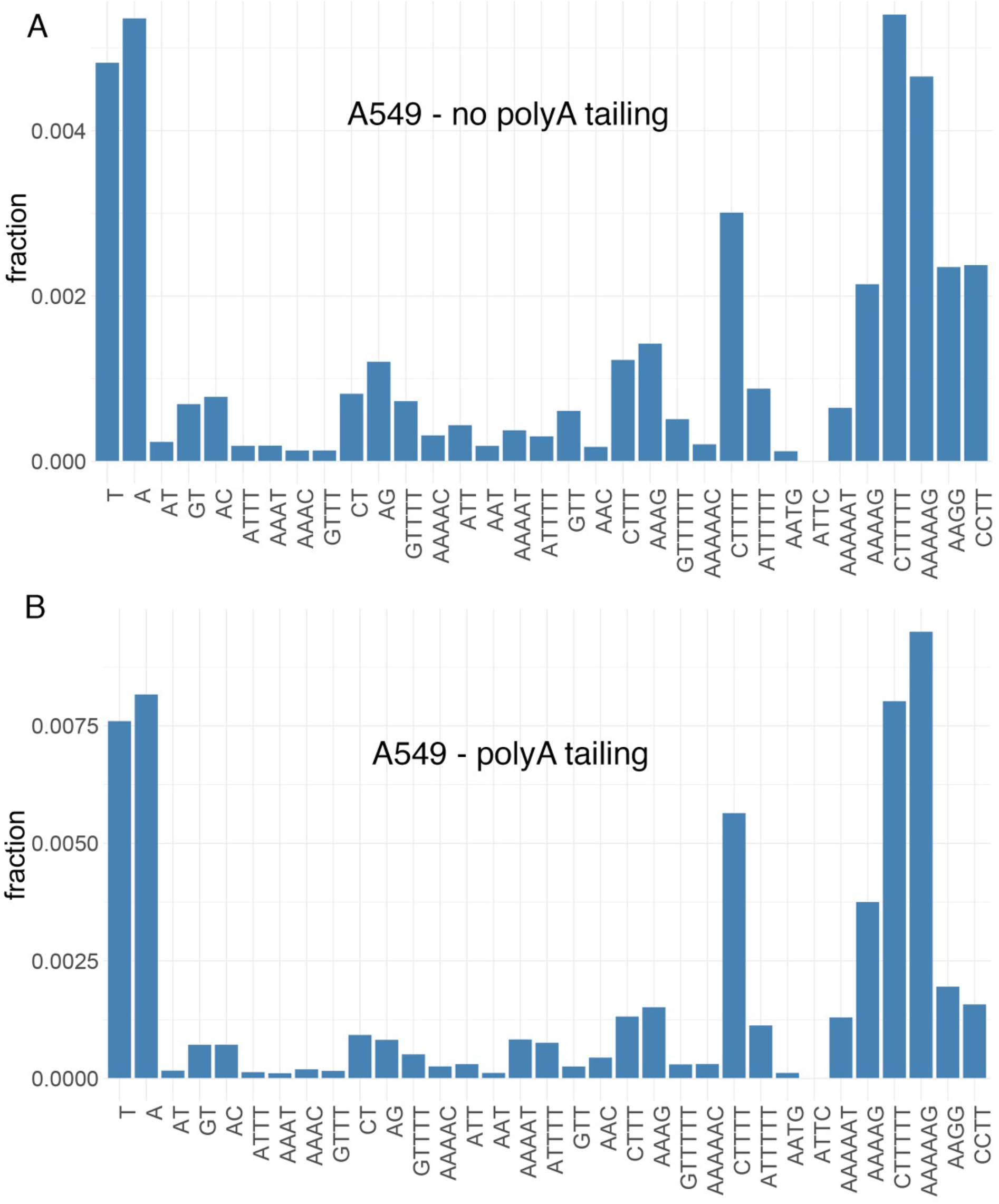
CTR-seq confirms the existence of transcription initiation at STRs. The fractions of STRs associated with at least one CTR-seq long read start site were computed for all STR classes considered in Figure 1B. RNAs were collected in A549 cells with (**B**) or without (**A**) polyA tailing before reverse transcription.

### Transcription at STRs exhibit specific features

We further looked at sub-cellular localization of STR-initiating transcripts and used CAGE sequencing data generated after cell fractionation (see Methods section). While the majority of CAGE tags, including those assigned to genes, are detected in both nucleus and cytoplasm, CAGE tags initiating at STRs are mostly detected in the nuclear compartment (Figure 4A). Functionally distinct RNA species were previously categorized by their transcriptional directionality^31^. We then sought to compute the directionality score, as defined in ^4^, for each STRs associated with CAGE signal (Figure 4B). Briefly, this score corresponds to the difference between the CAGE signal on the (+) strand and that on the (-) strand divided by their sum. A score equals to 1 or −1 indicates that transcription is strictly oriented towards the (+) or (-) strand respectively. A score close to 0 indicates that the transcription is balanced and that it occurs equally on the (+) and (-) orientation. As shown Figure 4B, some STR classes are associated with directional transcription either on the (+) (e.g. (*ATTT*)_*n*_, (*T*)_*n*_) or (-) (e.g. (*A*)_*n*_, (*ATG*)_*n*_) strand, while others are bidirectional and balanced ((*CGG*)_*n*_, (*CCG*)_*n*_). Note that the HipSTR catalog uses the (+) strand to define a DNA repeat motif. Therefore, (*A*)_*n*_ and (*T*)_*n*_ for instance do not correspond to the same loci. Transcription at (*A*)_*n*_, which is mostly detected on the (-) strand, does confirm the observation that transcription at (*T*)_*n*_ is mostly (+). The fact that transcription can be either directional or bidirectional depending on the STR class suggests that transcription at STRs is governed by different instructions, which are specific to STR classes.

**Figure 4.**
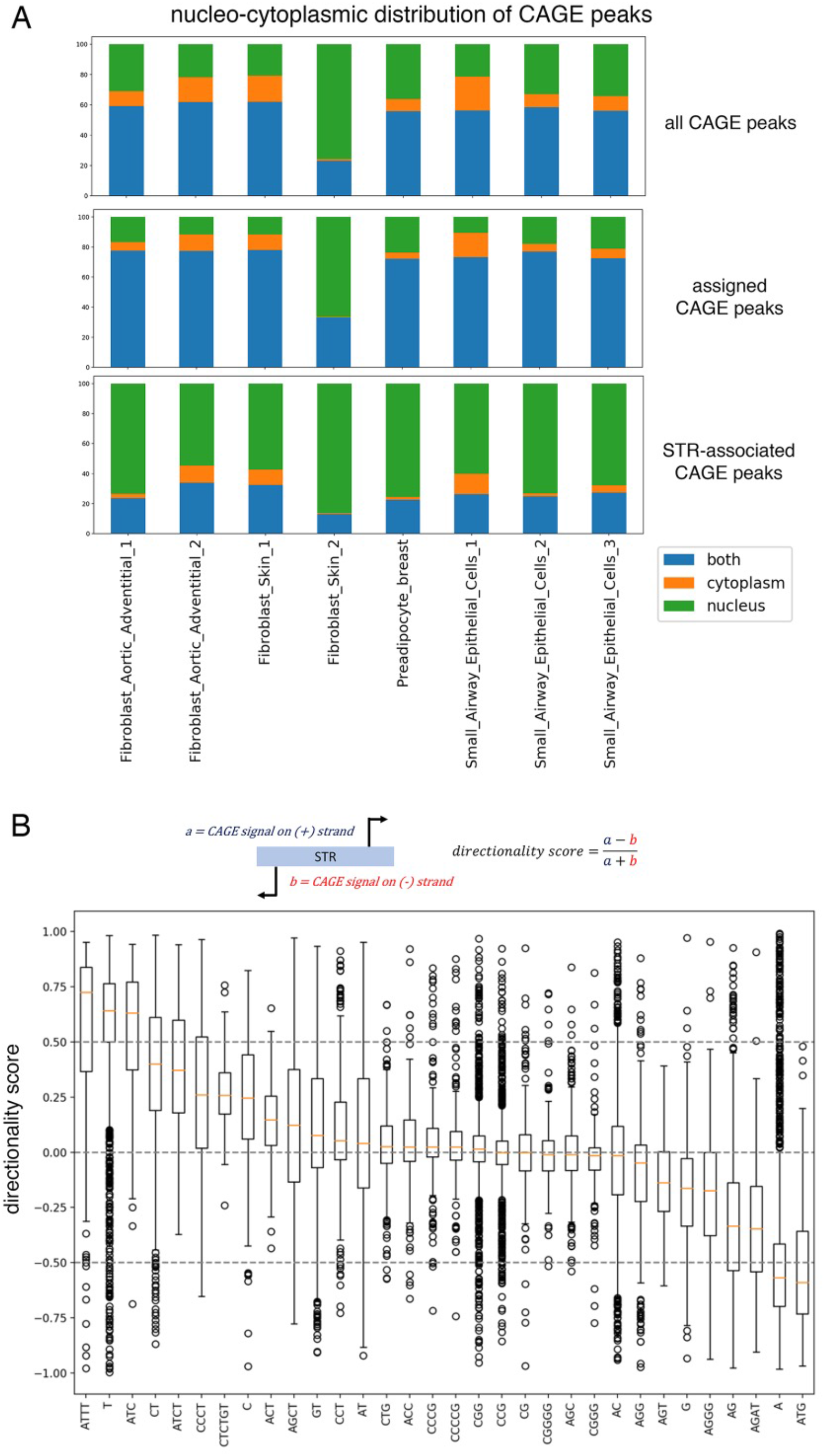
CAGE peaks at STRs exhibit specific features. **A.** STR-associated CAGE tags are preferentially detected in the nuclear compartment. For each indicated library (x-axis) and each CAGE peak, CAGE expression (TPM) was measured in nuclear and cytoplasmic fractions. Each CAGE peak was then assigned to nucleus (if only detected in the nucleus), cytoplasm (if only detected in the cytoplasm) or both compartments (if detected in both compartments). The number of CAGE peaks in each class is shown for each sample as a fraction of all detected CAGE peaks. The sample *Fibroblast Skin 2* likely represents a technical artifact. Analyses were conducted considering 201,802 FANTOM5 CAGE peaks (top), 54,001 CAGE peaks assigned to genes (middle), and 14,509 CAGE peaks associated with STRs (bottom). **B.** Boxplots of directionality scores for each STR classes with > 100 elements. A score of 0 means that the transcription is bidirectional and occurs on both strands. A score of 1 indicates that transcription occurs on the (+) strand, while −1 indicates transcription exclusively on the (-) strand.

### Probing transcription at STRs using a sequence-based deep learning model reveals that STR class are distinguishable

We further probed transcription at STRs using a machine learning approach. We used deep Convolutional Neural Network (CNN), which is able to successfully predict CAGE signal in large regions of the human genome ^32, 33^. This type of machine learning approches takes as input the DNA sequence directly, without the need to manually define predictive features before analysis. The first question that arose was then to determine the sequence to use as input.

We first sought to build a model common to all STR classes to predict the CAGE signal as computed in Figure 1D. Note that, because we used mean signal across CAGE libraires, our model is cell-type agnostic. This choice was motivated by the observation that the CAGE signal at STRs in each library is very sparse, thereby strongly reducing the prediction accuracy of our model. As input, we used sequences spanning 50bp around the 3’ end of each STR. Model architecture and constructions of the different sets used for learning are detailed in the Methods section and in Supplementary Figure S6. Source code is available at https://gite.lirmm.fr/ibc/deepSTR. The accuracy of our model was computed as Spearman correlation between the predicted and the observed CAGE signals. The performance of this global model was overall high (*ρ* ~ 0.72), indicating that transcription at STRs can indeed be predicted by sequence-level instructions. However, looking at the accuracy for each STR classes, we noticed drastic differences and the accuracy ranges from 0.66 to 0.81 depending on the STR class (Figure 5A). The global model is notably accurate for the most represented STR class (i.e. (*T*)_*n*_ with 766,747 elements) but performs less well in other STR classes. Differences in accuracies are not simply linked to the number of elements available for learning in each STR class. They rather suggest that, as proposed above (Figure 4B), transcription may be governed by instructions specific to each STR class.

**Figure 5.**
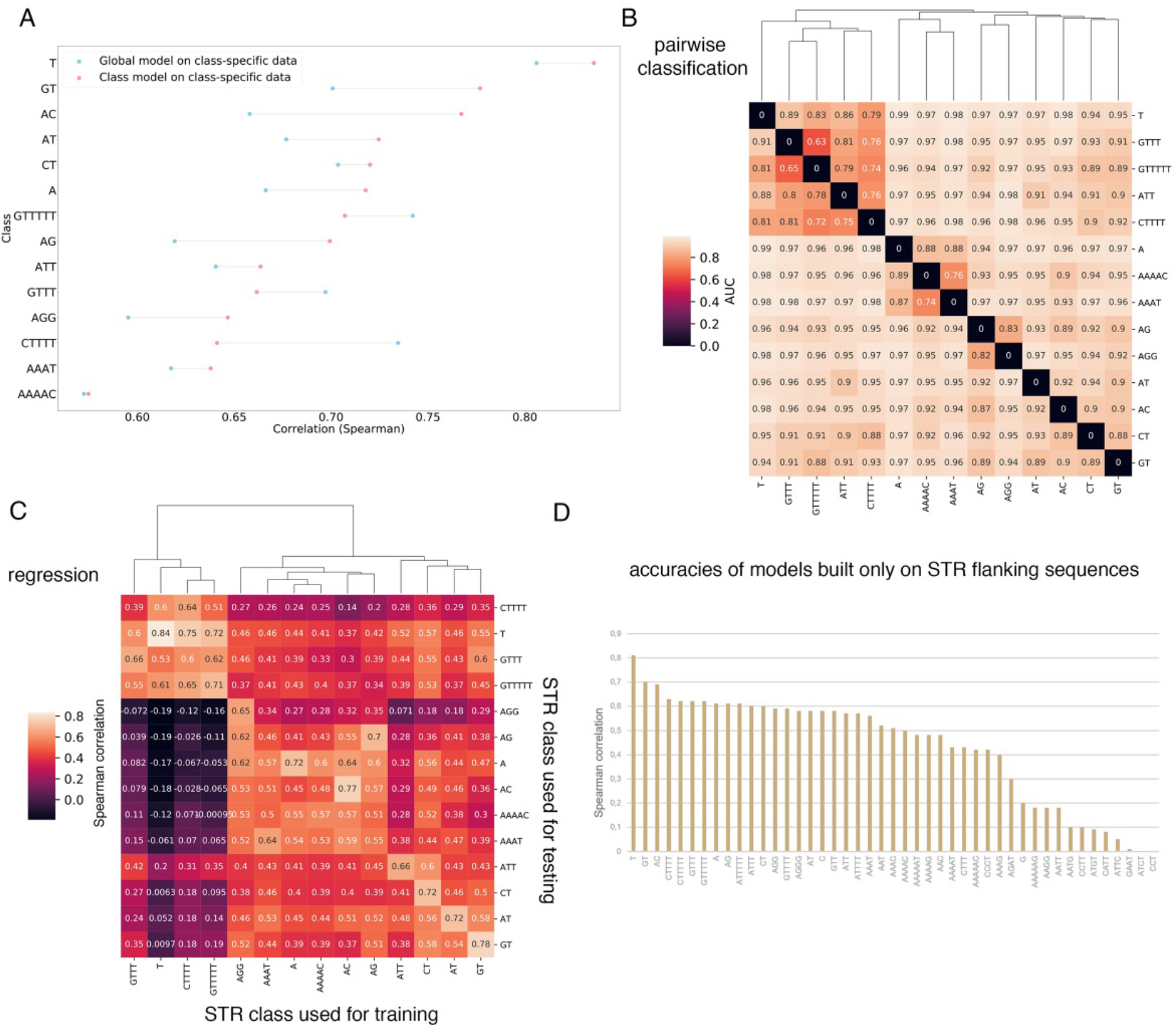
Probing STR sequences with CN models. **A.** Comparison of the accuracies of global vs. class-specific models. A model was learned on all STR sequences, irrespective of their class, and tested on each indicated STR classes (accuracies obtained in each case, as Spearman *ρ*, is shown as blue points). Distinct models were also learned for each indicated classes, without considering others (accuracies are shown in red). **B.** CNN-based pairwise classification of STRs using only STR flanking sequences (see Methods section). The pairs are defined by the line and the column of the matrix (i.e. the bottom left tile represents a classification task between T flanking sequences and A flanking sequences). The values displayed on the tiles correspond to AUCs measured on the test set with the model trained specifically for the task. Clustering was performed to group pairs of STRs according to AUCs. **C.** CNN performances to predict transcription at STRs evaluated as the Spearman correlation between predicted and observed CAGE signal. The heatmap represents the performance of one model learned on one STR class (rows) and tested either on the same or another class (columns). Clustering is also used to show which models are similar (high correlation) and which ones differ (low correlation). **D.** CNN models were learned on flanking sequences. The models use as input only the 50bp-long sequences flanking the STR, which is hidden and replaced by 9Ns (vectors of zeros in the one-hot encoded matrix). See Methods section.

### STR flanking sequences distinguish each STR class from others

It was previously shown that (*AC*)_*n*_ flanking sequences have evolved unusually to create specific nucleotide patterns ^34^. To determine if this specific pattern holds true for other STRs, we sought to classify STRs based only on their surrounding sequences. We trained a CNN model to classify pairs of STR classes (Supplementary Figure S6). To avoid any problem due to imprecise definition of STR boundaries, we masked the 7bases located downstream the STR 3’ ends (see Methods). In that case, model performance is evaluated by the Area Under the ROC (Receiver Operating Characteristics) curve (AUC, Figure 5B). The AUCs obtained in these pairwise classifications were very high (AUC > 0.7, Figure 5B), with the notable exceptions of (*T*)_*n*_ vs. (*GTTT*)_*n*_ ad (*T*)_*n*_ vs. (*GTTTTT*)_*n*_ (see below). Thus, STRs can be accurately distinguished, one from each other, using only flanking sequences, and strikingly, even in the case of complementary STRs, such as (*AC*)_*n*_ and (*GT*)_*n*_ (Figure 5B).

### Deep learning models unveil key role of STR flanking sequences

To further probe the sequence-level instructions for transcription at STRs, we decided to build a model for each STR class with > 5,000 elements (n = 47). Here, CNN is again used in a regression task to predict the CAGE signal as in Figure 5A. Sequences spanning 50bp around the 3’ end of each STR were used as input. Longer sequences were tested without improving the accuracy of the model (Supplementary Figure S7). These class-specific models achieved overall better performance than the global model tested on each STR class separately (Figure 5A). The only exceptions were classes composed of repetitions of T ((*GTTTTT*)_*n*_, (*GTTT*)_*n*_ and (*CTTTT*)_*n*_). In these cases, global and (*T*)_*n*_-specific models achieved better performance than specific model. These results have two explanations: (i) compared to (*T*)_*n*_, these classes have less occurrences (18,707 for (*GTTTTT*)_*n*_, 55,898 for (*GTTT*)_*n*_ and 15,433 for (*CTTTT*)_*n*_), making hard to learn models for these classes and (ii) the classification AUCs to distinguish (*GTTTTT*)_*n*_, (*GTTT*)_*n*_ or (*CTTTT*)_*n*_ from (*T*)_*n*_ were among the lowest observed (Figure 5B), suggesting the existence of common sequence features that can be used by global and (*T*)_*n*_-specific models. Overall, we estimated that these specific models were accurate for 14 STR classes (*ρ* > 0.68, Figure 5A).

We anticipated that class-specific models should not be equivalent and could not be interchangeable. We formally tested this hypothesis by measuring the accuracy of a model learned on one STR class and tested on another one (Figure 5C). We caution again the fact that the performance of a STR-specific model also depends on the number of sequences available for learning. As observed earlier, the best accuracy is obtained with (*T*)_*n*_, which are over-represented in our catalog. Overall, the performance of one model tested on another STR class drastically decreases (Figure 5C), formally demonstrating the existence of STR class-specific features for transcription prediction. We also noticed that several models achieved non negligible performance on other STR classes (Spearman *ρ* > 0.5, Figure 5C), implying that some features governing transcription at STRs are conserved between these STR classes. Thus, CNN models identified both common and specific features able to predict transcription at STRs.

Our results unveil the importance of STR flanking sequences. We then evaluated the contribution of the sole surrounding sequences in transcription prediction and built a model considering only these sequences (50bp upstream and downstream STR, masking the STR itself). These models were less accurate that the formers but accuracies were still high for several classes (Figure 5D), confirming that surrounding sequences contain instructions for transcription. The observed decrease in accuracies (Figure 5D) implies that the STR itself contains instructions, which are combined with features present in flanking regions to predict transcription. Remember that the CAGE signal predicted by our CNN models is normalized by the length of the STR (see above), which makes them unable to assess the contribution of STR length in transcription.

### Sequence-level instructions for STR transcription are conserved between human and mouse

To test whether transcription at STRs is biologically relevant, we relied on two criteria: conservation and association with diseases. First, we studied conservation in mouse.

The abundance of STR classes globally differs between mouse and human genomes, except for highly abundant classes (e.g. (*T*)_*n*_, (*A*)_*n*_, (*AC*)_*n*_, (*GT*)_*n*_ or (*AT*)_*n*_, compare Figure 1B and Figure 6A). We applied the strategy used in human to compute the CAGE signal (as mean raw tag count in STR ± 5bp divided by STR length + 10bp) in mouse using 397 CAGE libraries (Figure 6B). As observed in human, several STR classes were associated with CAGE signal. This signal appears lower than in human (compare Figure 1D and Figure 6B). This might be due to the fact that mouse CAGE data are small-scaled in terms of number of reads mapped and diversity in CAGE libraries, compared to human CAGE data ^2^, making the mouse CAGE signal at STRs probably less reliable that the human one.

**Figure 6.**
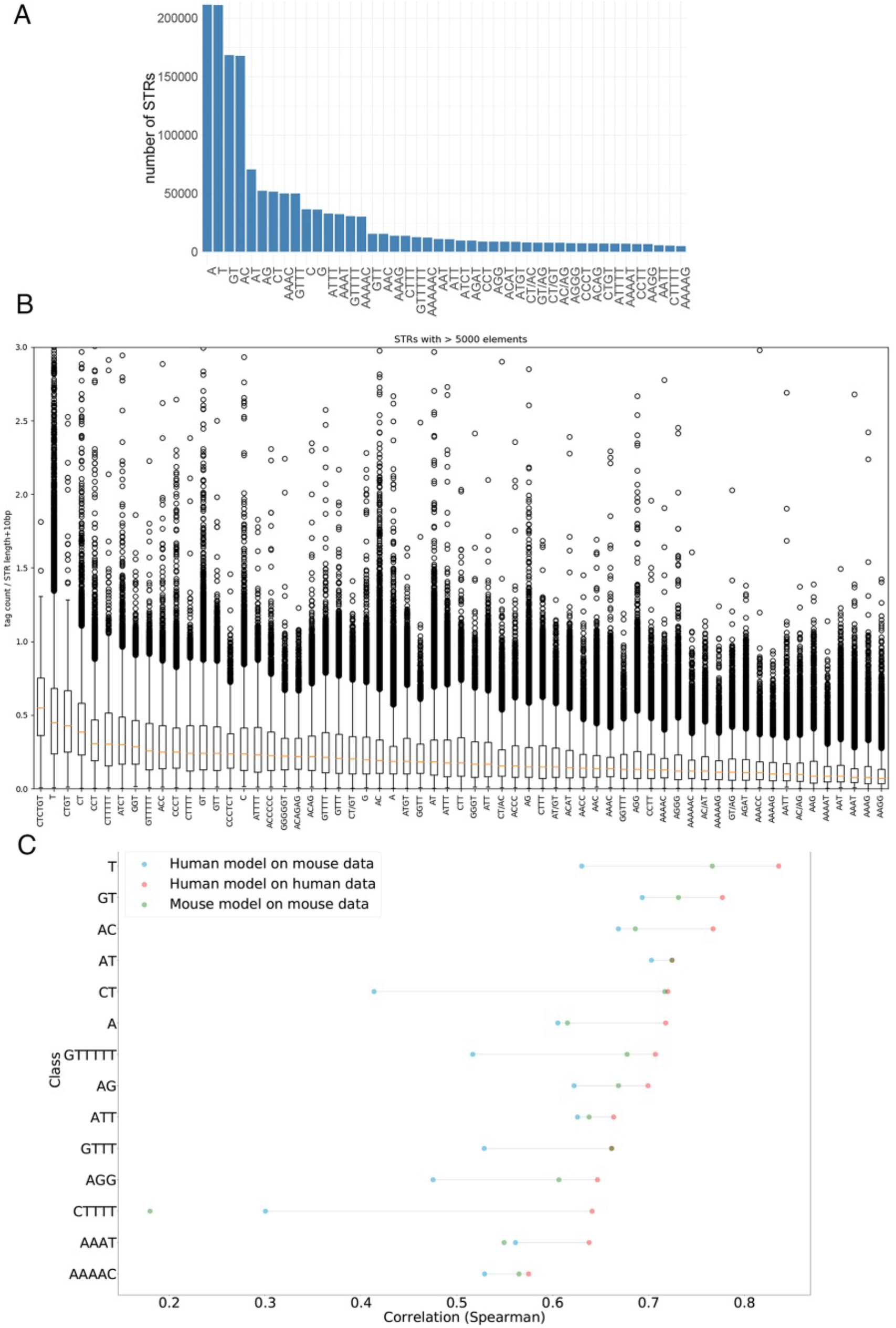
STR transcription in mouse. **A.** Number of mouse STRs per class. For sake of clarity, only STR classes with > 5,000 loci are shown. **B.** CAGE signal at mouse STR classes with > 5,000 loci. CAGE signal was computed as in Figure 1D. **C.** Testing the accuracy of CNN models built in human and tested in mouse for different STR classes. Performance of the model is assessed by computing the Spearman *ρ* between CAGE signal observed in mouse and that predicted by a model learned in human (blue dots), CAGE signal observed in mouse and that predicted by a model learned in mouse (green dots) and CAGE signal observed in human and that predicted by a model learned in human (green dots).

We nonetheless tested the correlation of the human and mouse CAGE signals at orthologous STRs. Orthologous STRs were identified converting the mouse STR coordinates into human coordinates with the UCSC liftover tool (see Methods). We intersected the coordinates of human STRs with that of orthologous mouse STRs and computed the Pearson correlation between the CAGE signal observed in human and that observed in mouse on the same strand (n = 18,072). In that case, Pearson’s r reaches ~ 0.87 (Spearman *ρ* ~ 0.51), suggesting that transcription at STRs is indeed conserved between mouse and human. As expected, no correlation was observed (r < 0.01) when randomly shuffling one of the two vectors or when correlating the signals of 18,072 randomly chosen mouse and human STRs.

We then built a CNN model to predict CAGE signal at mouse STR classes corresponding to the 14 classes well predicted in human (Figure 6C, green dots). The performances of the models ranged from ~ 0.4 to ~ 0.8, demonstrating that, as observed for human STRs, transcription at several mouse STR classes can be predicted by sequence-level instructions. Notable exception is (*CTTTT*)_*n*_ with Spearman *ρ* < 0.2 (see below). The mouse models were overall less accurate than human models (Figure 6C, compare red and green dots), likely due to differences in the quality of the CAGE signal (i.e. predicted variable), as mentioned above.

We then tested whether the sequence features able to predict STR transcription were conserved between mouse and human. We specifically tested the performances of models learned in one species and tested on another one (Figure 6C and Supplementary Figure S8). For all STR classes tested, the Spearman correlation between the signal predicted by the human model and the observed mouse signal was > 0.4 (Figure 6C), implying that several features are conserved between human and mouse. For some classes (e.g. (*A*)_*n*_, (*AC*)_*n*_, (*AAAT*)_*n*_), the human and mouse models even appeared equally efficient in predicting transcription in mouse (Figure 6C, green and blue dots are close), indicative of a strong conservation of predictive features. For other classes (e.g. (*CT*)_*n*_, (*AGG*)_*n*_), the performance of the human model was lower than that obtained with the mouse model when tested on mouse data (Figure 6C, green and blue dots are distant). Thus, specific features also exist in mouse that were not learned in human sequences. Likewise, human specific features also exist (Supplementary Figure S8). Strikingly, in the case of (*CTTTT*)_*n*_, the human model was even able to predict mouse CAGE signal, when the mouse model was not (Figure 6C). This effect is likely due to the number of examples, which is higher in human (n = 15,433) than in mouse (n = 10,494). Overall, we concluded that several features predictive of transcription at STRs are conserved between human and mouse and that the level of conservation also varies depending on STR classes.

### Clinically relevant genetic variants are predicted to impact STR transcription

Second, we evaluated the potential implication of STR transcription in human diseases and used the ClinVar database, which lists medically important variants ^35^. We found that 34,578 STRs harbour at least one ClinVar variants in a window encompassing STR ± 50bp. Strikingly, these STRs are associated with high CAGE signal compared to STRs without variants (n = 3,068,280, Figure 7A), indicative of potential biological and clinical relevance for STR transcription. The clinical significance of the variants, as defined in theClinVar database ^35^, does not appear directly linked to the transcription rate of STRs (Supplementary Figure S9). Likewise, several diseases were found enriched comparing variant fractions located at transcribed STRs (Fisher’s exact test < 5e-3, Supplementary Table S2) but no clear association with a specific clinical trait was noticed.

**Figure 7.**
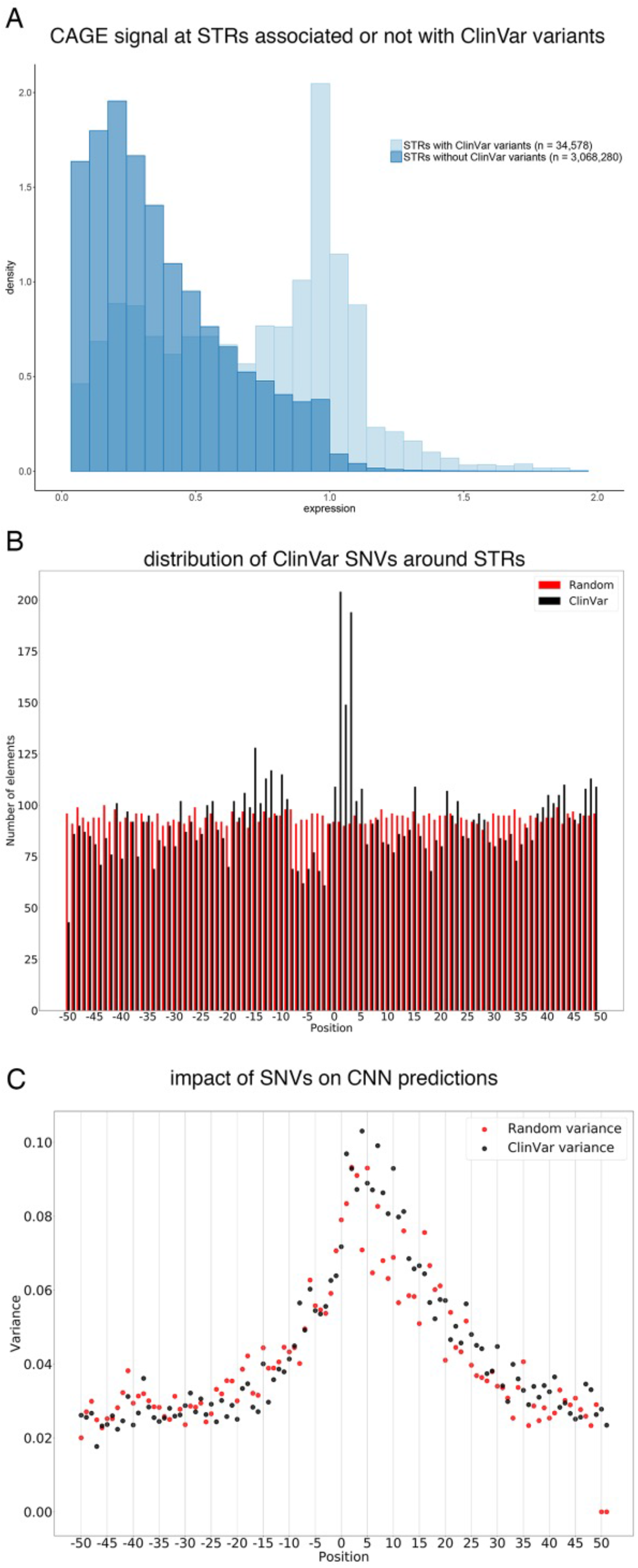
Assessing the impact of ClinVar variants on STR transcription. **A.** CAGE signal distribution of STRs associated (light blue) or not (dark blue) with at least one ClinVar variant. The number of STRs considered in each case is indicated in bracket. **B.** Distribution of ClinVar (black) and random (red) variants around STR 3’end. The number of variants and their position relative to STR 3’end (position 0) are indicated on the y-axis and x-axis respectively. **C.** Impact of the changes induced by ClinVar (black) and random (red) variants in CNN predictions. Predictions are made on the hg19 reference sequence and on a mutated sequence, containing the genetic variants. Changes are then computed as the difference between these two predictions (reference – mutated, Supplementary Figure S10) and their impact is measured as their variance at each position around STR 3’ end (x-axis). To keep sequences aligned, only single nucleotide variants were considered.

Moreover, there is a significant proportion of ClinVar variants located in the immediate vicinity of the STR (Figure 7B), indicating that these positions may often be more important from a clinical point of view.

We then used a perturbation-based approach ^36^ and randomly created *in silico* mutations to identify key positions of the models (Figure 7B and see Methods section). Random variations were directly introduced into STR sequences and predictions were made on these mutated sequences using the CNN model specific of the STR class considered. The impact of the variation was then assessed as the difference between the predictions obtained with mutated and reference sequences. Same analyses were performed with ClinVar variants (Supplementary Figure S10). Key positions were defined as positions, which, when mutated, have a strong impact on the prediction changes (i.e. high variance), being either positive or negative. As shown Figure 7C, for both random and ClinVar variants, the most important positions appeared located around STR 3’end and their distribution is skewed towards the sense orientation of the transcripts. Hence, the most important positions identified by our models correspond to positions with high occurrences of ClinVar variants (Figures 7B and 7C). This indicated that our models have learned key positions to predict transcription at STRs and that these positions correspond precisely to genetic variants linked to human diseases. Note that a similar distribution is observed for ClinVar variants around assigned CAGEs but not around all identified CAGE peaks (Supplementary Figure S11).

## Discussion

The human genome is scattered with repetitive sequences, and the vast majority of the genome is supposed to be transcribed ^1, 5, 6^. The human transcriptome should then contain a large portion of RNAs derived from repetitive elements ^17^, including STRs. Previous works have already shown that several classes of repetitive elements can be transcribed, from retrotransposons ^37^ to DNA satellites^19, 20^. Here, using FANTOM CAGE data ^2^, we provide evidence that a significant fraction of STRs, from distinct classes, are also transcribed in both human and mouse, a process which further complexifies STR polymorphism.

RNA species can be functionally categorized according to transcriptional directionality ^31^. In the case of STRs, transcription directionality appears to depend on the STR class (Figure 4B). It is thus likely that RNAs initiating at STRs fulfil distincts function and many hypotheses could be proposed. For instance, 10,727 CAGE peaks mapped at STRs correspond to TSSs of FANTOM CAT transcripts (Supplementary Table S1), extending the findings made by by Bertuzzi *et al*. in the case of minisatellites and NPRL3 gene ^20^ to STRs. Many RNAs initiating at STRs may also correspond to non-coding RNAs, as for instance enhancer RNAs (Supplementary Table S1). As could have been anticipated given the distinction of enhancers and promoters based on CpG dinucleotide ^38^, FANTOM CAT transcripts mostly initiate at GC-rich STRs, while enhancer RNAs more often correspond to A/T-rich STRs (Supplementary Table S1). Another possible function is provided by (*T*)_*n*_, which are over-represented in eukaryotic genomes^39^ and have been shown to act as promoter elements by depleting repressive nucleosomes ^40^. As a consequence, (*T*)_*n*_ can increase transcription of reporter genes in similar levels to TF binding sites ^41^. The findings that (*A*)_*n*_ and (*T*)_*n*_ represent distinct directional signals for nucleosome removal^42^, are very well compatible with differences observed in flanking sequences (Figure 5B) and directional transcription (Figure 4B), both able to create asymmetry at (*A*)_*n*_ and (*T*)_*n*_. Besides, we show that most CAGE tags initiating at STRs remain nuclear (Figure 4A). This observation suggests that, similar to other repeat-initiating RNAs ^43, 44^, STR-initiating RNAs could also play a role in DNA topology ^44, 45^. At this stage, it remains to clarify whether STR-associated RNAs or the act of transcription *per se* is functionally important ^10^. Dedicated experiments are now required to formally identify the biological functions linked to the transcription of each STR class. These experiments are all the more warranted as STR transcription is associated with clinically relevant genomic variations (Figure 7).

One key finding of our study is the discovery that STR flanking sequences are not inert but rather contain important features that play critical roles in their biology, as previously suspected^34^. These results call for the development of novel methods able to take these sequences into account in order to revisit STR mapping/genotyping and integrate SNVs located at STR vicinity. These methods should have broad applications in various fields of research and medicine, from forensic medicine to population genetics for instance. STR length variations have notably been shown to influence gene expression and, similar to expression quantitative trait loci (eQTLs), several expression STR (eSTR) have been identified ^46, 47^. Their exact mode of action still remain largely elusive but, the majority of eSTRs appears to act by global mechanisms, in a tissue-agnostic manner ^46^. Interestingly, some eSTRs have strand-specific effects ^46^, which is again compatible with the possible sources of asymmetry unveiled by our study (i.e. flanking sequences and directional transcription). Using transcription level of STRs, as predicted by our CNN models for instance, coupled with length variations ^46, 47^, may help to take into account the impact of genetic variants located in sequences surrounding STRs, and to refine eSTR computations.

There are still several ways to improve our CNN models. Notably, to avoid any bias linked to CAGE noise signal observed along STRs, we decided to predict a signal normalized by the STR length. Therefore, our models do not allow to properly assess the contribution of STR length in transcription, although it clearly represents the most studied features of STRs ^21, 46, 47^. Note that simply increasing the quality of the reads considered (using Q20 instead of Q3 filter) yielded sparse data and decreased the performance of our model. A new computation of the CAGE signal aimed at removing ‘noise’ at STRs could be developed. This may also help develop tissue-specific CNN models, which will only use CAGE data ^33^. Besides, the same architecture was used for all STR classes while achieving different accuracies (Figures 5A and C). These results cannot be merely explained by the varying numbers of STR sequences available for training because swapping the models for training and testing demonstrated the existence of STR class-specific features predictive of transcription (Figure 5C). Rather, the chosen architecture may not be optimal for all STRs, as illustrated by the design of a global model with overall good performance, but very distinct accuracies depending on the STR class (Figure 5A). Our CNN architecture was initially optimized on the (*T*)_*n*_ class, which represents the most abundant class (n = 766,747). Because each STR class harbour sequence specificities including in flanking sequences, hyperparameters, such as convolutional filter sizes, their number and/or max-pooling, could be adapted to each STR class. These hyperparameters have indeed already been shown to influence the results of CNN models as well as their interpretation ^48^.

More broadly, the same rationale could be applied to other methods aimed at predicting CAGE signal along the genome ^33^, distinguishing biological entities (genes, enhancers, …), genomic segments ^49, 50^ and/or isochores ^51^ based on their sequence features. Building a general model increases the risk of designing a model suited for the most represented elements, not for the others. Notably, promoters and enhancers can be distinguished by different CpG content, the presence of polyA signal and of 5’ splice sites^38^, as well as different transcription factor combinations ^3, 52^. It is therefore likely that the same filters will not apply similarly to predict transcription in both cases and that one may want to develop a specific model for each of these entities to increase the accuracy of the predictions.

The prediction of transcription initiation based solely on sequence features has long been studied, especially using CAGE data ^53, 54^. The high accuracy achieved by CNN models for this task, as illustrated in this study or in ^32, 33^, as well as the development of methods aimed at interpreting this type of statistical models ^36, 48, 55, 56^, will certainly accelerate the achievement of this goal, which becomes more than ever ‘a realistic short-term objective rather than a distant aspiration’^54^.

## Methods

### Data and bioinformatic analyses

The bedtools window ^57^ was used to look for CAGE peaks (http://fantom.gsc.riken.jp/5/datafiles/phase1.3/extra/CAGE_peaks/hg19.cage_peak_coord_permissive.bed.gz) at STRs ± 5bp (https://github.com/HipSTR-Tool/HipSTR-references/raw/master/human/hg19.hipstr_reference.bed.gz) as followed:

~~~
windowBed -w 5 -a hg19.hipstr_reference.bed -b hg19.cage_peak_coord_permissive
   .bed
~~~

Similar analyses were performed using mouse STR catalog: https://github.com/HipSTR-Tool/HipSTR-references/blob/master/mouse/mm10.hipstr_reference.bed.gz) liftovered to mm9 using UCSC liftover tool^58^ as:

~~~
liftover mm10.hipstr_reference.bed mm10ToMm9.over.chain.gz mm9.
   hipstr_reference.bed unlifted.bed
~~~

To compute the CAGE signal, we used raw tag count along the genome with a 1bp binning and Q3 quality mapping filter. At each position of the genome, the mean tag count across 988 librairies for human and 387 for mouse was computed. The values obtained at each position of a window encompassing the STR± 5bp were then summed and normalized (i.e. divided by the STR length + 10 bp) to limit the impact of CAGE noise signal observed along STRs. CAGE signals at human and mouse STRs are available at https://gite.lirmm.fr/ibc/deepSTR as respectively hg19.hipstr_reference.cage.bed and mm9.hipstr_reference.cage.bed (The CAGE signal is indicated in the 5th column). The fasta files (500 bp around STR 3’end) used to build our models are also available at the same location as hg19.hipstr_reference.cage.500bp.around3end.fa and mm9.hipstr_reference.cage.500bp.around3end.fa

The bedtools intersect^57^ was used to distinguish intra- and intergenic STRs, intersecting their coordinates with that of the FANTOM gene annotation https://fantom.gsc.riken.jp/5/suppl/Hon_et_al_2016/data/assembly/lv3_robust/FANTOM_CAT.lv3_robust.bed.gz.

FANTOM CAT robust transcripts coordinates can be found at http://fantom.gsc.riken.jp/5/suppl/Hon_et_al_2016/data/assembly/lv3_robust/FANTOM_CAT.lv3_robust.gtf.gz and that of FANTOM enhancers at https://fantom.gsc.riken.jp/5/datafiles/latest/extra/Enhancers/human_permissive_enhancers_phase_1_and_2.bed.gz

ENCODE RNAPII ChIP-seq bed files can be donwloaded at http://hgdownload.cse.ucsc.edu/goldenpath/hg19/encodeDCC/wgEncodeAwgTfbsUniform/wgEncodeAwgTfbsHaibGm12878Pol2Pcr2xUniPk.narrowPeak.gz, http://hgdownload.cse.ucsc.edu/goldenpath/hg19/encodeDCC/wgEncodeAwgTfbsHaibH1hescPol2V0416102UniPk.narrowPeak.gz, http://hgdownload.cse.ucsc.edu/goldenpath/hg19/encodeDCC/wgEncodeAwgTfbsHaibHelas3Pol2Pcr1xUniPk.narrowPeak.gz and http://hgdownload.cse.ucsc.edu/goldenpath/hg19/encodeDCC/wgEncodeAwgTfbsHaibK562Pol2V0416101UniPk.narrowPeak.gz.

Expression data used to determine the nucleo-cytoplasmic distribution of CAGE peaks can be found at http://fantom.gsc.riken.jp/5/datafiles/latest/extra/CAGE_peaks/hg19.cage_peak_phase1and2combined_tpm_ann.osc.txt.gz.

Orthologous STRs were identified using UCSC liftover tool^58^ and the mm9ToHg19.over.chain.gz file.

### Evaluating mismatched G bias at Illumina 5’end CAGE reads

Comparison between Heliscope vs. Illumina CAGE sequencing was performed as in de Rie *et al*.^30^. Briefly, ENCODE CAGE data were downloaded as bam file (http://hgdownload.cse.ucsc.edu/goldenpath/hg19/encodeDCC/wgEncodeRikenCage/*NucleusPap*) and converted into bed file using samtools view ^59^ and unix awk as follow:

~~~
samtools view file.bam | **awk** ‘{FS=“\t”}BEGIN{OFS=“\t”}{**if**($2==“0”) print $3,
   $4-1,$4,$10,$13,”+”; **else if**($2==“16”) print $3,$4-1,$4,$10,$13,”-”}’ >
   file.bed
~~~

The bedtools intersect ^57^ was further used to identify all CAGE tags mapped at a given position. The unix awk command was used to count the number and type of mismatches as follow:

~~~
intersectBed -a positions_of_interest.bed -b file.bed -wa -wb -s |
**awk** ‘{**if**(substr($11,1,6)==“MD:Z:0” && $6==“+”) print substr($10,1,1)}’ | grep
   -c “N”
~~~

with N = {A, C, G or T}, positions of _interest.bed being coordinates of CAGE peaks assigned to genes, or that located at pre-miRNA 3’ ends, or peaks associated with STR. The file.bed corresponds to the Illumina CAGE tag coordinates.

Absence of mismatch focusing on the plus strand were counted as:

~~~
intersectBed -a positions_of_interest.bed -b file.bed -wa -wb -s |
**awk** ‘{**if**(substr($11,1,6)!=“MD:Z:0” && $6==“+”) print $0}’ |wc -l
~~~

As a control, we used the 3’ end of the pre-miRNAs, which were defined, as in de Rie *et al*.^30^, as the 3’ nucleotide of the mature miRNA on the 3’ arm of the pre-miRNA (miRBase V21, ftp://mirbase.org/pub/mirbase/21/genomes/hsa.gff3), the expected Drosha cleavage site being immediately downstream of this nucleotide (pre-miR end + 1 base).

### Cap-Trapping MinION sequencing

A549 cells were grown in Dulbeccos modified Eagle medium (DMEM) supplemented with 10% fetal bovine serum (FBS). A549 cells were washed with PBS. The RNAs were isolated by using RNeasy kit (QIAGEN). The poly-A tail addition to A549 total RNA was carried out by poly-A polymerase. (PAPed RNA) The cDNA synthesis was carried out by using 5 g of total RNA or 1 g of PAPed RNA with RT primer (5-TTTTTTTTUUUTTTTTVN-3) by PrimeScript II Reverse Transcriptase (TaKaRa Bio). The full-length cDNAs were selected by Cap Trapper method ^60^. After the ligation of 5 linker, cDNAs were treated with USER enzyme to shorten the poly-T derived from RT primer. After SAP treatment, 3 linker was ligated to the cDNAs. The linkers used in the library preparation were prepared by annealing using these oligos https://pubmed.ncbi.nlm.nih.gov/32124327/ with following oligos:

5’ linker GN5 up: 5- GTGGTAUCAACGCAGAGUACGNNNNN -P-3’
5’ linker N6 up: 5- GTGGTAUCAACGCAGAGUACNNNNNN -P-3’
5’ linker down: 5’-P- GTACTCTGCGTTGATACCAC-P-3’
3’ linker up: 5’-AAAAABBBBBBBBGCAUCGCUGTCTCUTAUACACAUCUCCGAGCCCACGAGAC -P-3
3’ linker down: 5’- GTCTCGTGGGCTCGGAGATGTGTATAAGAGACAGCGATGC -3’

As for the 3’ linker, after annealing step, the UMI complemental region (BBBBBBBB) was filled with Phusion High-Fidelity DNA polymerase (NEB) and dVTPs (dATP/dGTP/dCTP) instead of dNTPs. The 2nd strand was synthesized using 2nd primer (5-TCGTCGGCAGCGTCAGATGTGTATAAGAGACAGNNNNNNNNGTGGTATCAACGCAGAGTAC -3) with KAPA HiFi HS mix (KAPA Biosystems). The double stranded cDNAs were amplified using Illumina adapter-specific primers and LongAmp Taq DNA polymerase (NEB). After 16 cycles of PCR (8?minutes for elongation time), amplified cDNAs were purified with equal volume of AMPure XP beads (Beckmann Coulter). Purified cDNAs were subjected to Nanopore sequencing library following to manufacturers 1D ligation sequencing protocol (version NBE_9006_v103_revO_21Dec2016).

Nanopore libraries were sequenced by MinION Mk1b with R9.4 flowcell. Sequence data was generated by MinKNOW 1.7.14. Basecalling was processed by Albacore v2.1.0 basecaller software provided by Oxford Nanopore Technologies to generate fastq files from FAST5 files. To preparing clean reads from fastq files, adopter sequence was trimmed by Porechop v0.2.3.

Detailed protocol is provided as Supplementary Methods.

Data were deposited on DNA Data Bank of Japan Sequencing Read Archive (accession number: DRA010491).

The mapping computational pipeline used a prototype of primer-chop available at https://gitlab.com/mcfrith/primer-chop. The precise methods and command lines are provided as Supplementary Methods. Data were first mapped on hg38 reference genome and liftovered to hg19 for analyses.

### Directionality score

We collected CAGE signal at each STR of the HipSTR catalog (see above). When a signal was detected on both (+) and (-) strands, we computed the directionality score for each STR using the following formula:

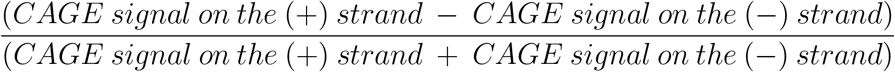

The CAGE signal was computed as explained above. A score equals to 1 or −1 indicates that transcription is strictly oriented towards the (+) or (-) strand respectively. A score close to 0 indicates that the transcription is balanced and that it occurs equally on the (+) and (-) orientation.

### Convolutional Neural Network

CNN architecture is described in Supplementary Figure S6. Sequences spanning 50bp around the 3’ end of each STR were used as input. Longer sequences were tested without improving the accuracy of the model (Supplementary Figure S7). Note that only 89,189 STRs (out of 1,620,030, ~ 5.5%) are longer than 50bp and, only in these few cases, the sequence located upstream STR 3’ end only corresponds to the STR itself. The parameters of the model (such as number of layers, number of neurons, optimizer, activation functions…) were determined by brut force algorithms. The model is implemented in PyTorch. the source code of the model, alongside scripts and Jupyter notebooks are available at https://gite.lirmm.fr/ibc/deepSTR.

In order to minimize overfitting, droupout is added to the fully connected layers (probability of droupout = 0.30). The training pipeline is described in Supplementary Figure S6: we separate training, testing and validation datasets prior to model training, and these sets are stored on disk. This allows us to perform analysis on data that the models have never seen. During training, we stop the training once the loss function calculated on the validation set drops for 5 consecutive epochs (early stopping). Note that relatively good performances on mouse datasets (Figure 6C) show that the model generalizes well to unknown data.

### Classification

The CNN model can also be set up for a classification task (Figure 5B). In that case, the only difference with the previous model is the last neuron in the last fully connected layer. The classifier CNN uses the same training method. The data are also prepared by separate scripts before training is done and stored on disk. All analyses resulting from classification are performed on the test sets to avoid optimistic bias in accuracy estimation. Note that 7bp were masked after STR 3’ end and replaced by Ns because we noticed that this window can contain bases corresponding to the DNA repeated elements, a feature that can easily be learned by a CNN. The sequence used as input are centered around STR 3’ end and consist of 50bp-long upstream sequence + 9 Ns, which mask the STR itself + 7 Ns + 43bp-long downstream sequence (total length = 109bp).

### Model swaps between human STRs classes

After models are trained on all STR classes, their weights are stored in a .pt file (following the PyTorch convention). Predictions were then computed on all test sets with all models.

### Predicting impact of ClinVar variants

ClinVar vcf file was downloaded January 8th 2019 (ftp://ftp.ncbi.nlm.nih.gov/pub/clinvar/) and then converted into bed file. We looked for STRs associated withClinVar variants (Figure 7A) using bedtools window^57^ as follows:

~~~
 bedtools window -w 50 -a clinvar_mutation.bed -b str_coordinates.bed
~~~

Variants were directly introduced into STR sequences (± 50bp) using Biopython ^61^ library and the *seq.tomutable()* function. To keep sequences aligned, we only considered single nucleotide variants (SNVs). The CNN model developed previously was then used to predict the CAGE signal of the initial and mutated sequences. The change was computed by the difference between the prediction obtained with mutated sequence and that obtained with the reference sequence.

To insert random variations (Figure 7B and 7C), we created a mutation position map, which follows a uniform distribution (each position has an equal probability of receiving a mutation). Then, we took sequences in the database and mutated them one by one at a position taken from the mutation map. All possible mutations at the chosen position have an equal probability of occurrence (see Figure 7B).

## Acknowledgements

We thank Cédric Notredame, Anthony Mathelier, Oriol Fornes Crespo, Philip Richmond, Jean-Christophe Andrau, Diego Garrido Martin, Dimitri D. Pervouchine, Roderic Guigo, Charles Plessy and Chung Hon for their help in analyzing the data and for insightful suggestions. We also thank Takahiro Arakawa for the preparation and provision of cell culture samples. We are indebted to the researchers around the globe who generated experimental data and made them freely available. C-H.L. is grateful to Marc Piechaczyk and Edouard Bertrand for continued support.

## Author contributions

C.B., M.S., M.G., C.M. W.W.W., M.d.H, L.B and C-H.L. analyzed and interpreted data. M.S. and M.G. developed CNN models and studied the impact of ClinVar variants. J.R., Y.H., A.H., H.S., S.N., I.M. generated CAGE data used in this study. M.d.H., J.S. and C-H.L. generated Zenbu tracks. M.d.H and C-H.L. studied G bias at ENCODE read 5’ ends. M.T., M. M., M. K-I., S. N., S. N.,T. K., H. N., M.F. developed CTR-seq and generated data used in this study. Y.H., P.C., C.C., W.W.W, L.B. and C-H.L acquired fundings. C-H.L. wrote the manuscript. All authors have read and approved the manuscript.

## Correspondence

Correspondence and requests for materials should be addressed to L.B. and C-H.L..

## Supplementary Methods: Cap Trap and MinION sequencing

### 1. Addition of polyA tail

The polyA tail addition was carried out by using 8 ug of totalRNA in 14.5ul of water, 2.0ul of 10x PolyA polymerase buffer (NEB), 2.0ul of 10mM ATP (NEB), 1.0ul of RNaseOUT (Invitrogen) and 0.5ul of PolyA polymeras(5 U/ul). We incubated this reaction mix at 37C for 15m, then put the tube on ice. After polyA polymerase reaction, polyA-tailed totalRNA (PAPed RNA) was purified with Agencourt RNAClean XP kit (Beckman coulter) according to the manufacturer’s instructions and eluted in 40ul of water.

### 2. Reverse Transcription

We put 5ul each of PAPed RNA into 8 wells.

The cDNA synthesis was carried out by using 5ug of total RNA or 1ug of PAPed RNA in 5ul of water and 0.5ul of 100uM RT primer (5’- TTTTTTTTUUUTTTTTVN -3’) by PrimeScript II Reverse Transcriptase(TaKaRa). We heated RNA and primer at 65C for 5min and then placed them on ice. Then we added the reaction mixture, 4ul of 5x PrimeScript II buffer, 4ul of water, 1ul of RNaseOUT and 1ul of PrimeScript II, followed by reverse transcription in a thermal cycler: 42C for 60min, then chilled at 4C.

After the reaction, the cDNA/RNA hybrids were purified with Agencourt RNAClean XP.

### 3. Oxidation / Biotinylation

To oxidize the diol residue of Cap structure, 40ul of purified cDNA/RNA hybrids were mixed with 2ul of 1M NaOAc (pH4.5) and 2ul of 250mM NaIO4 (Sigma-Aldrich) and incubated on ice for 5min in dark. To stop the reaction, the oxidized cDNA/RNA hybrids were mixed with 16ul of 1M Tris-HCl(pH8.5). The sample was purified with RNAClean XP. Four ul of 1M NaOAc (pH6.2) and 4ul of 100mM Biotin (long arm) hydrazide (Vector Laboratories) in DMSO were added and the reaction mixture were incubated at 40C for 30min. After the incubation, the biotinylated sample was purified with RNAClean XP. Finally, single-strand RNA regions which were not protected by a complementary first-strand cDNA strand were digested using RNaseONE(Promega) by addition of 4.5ul of 10×RNaseI buffer and 0.5ul of RNaseONE and incubation at 37C for 30min. The reaction mixture were purified with RNAClean XP.

### 4. CapTrap

Thirty microliters of Dynabeads M-270 Streptavidin beads slurry (ThermoFisher Scientific) was washed with 30ul of LiCl binding buffer (7M LiCl, 10mM Tris-HCl (pH7.5), 0.1% Tween20, 2mM EDTA (pH8.0)) twice and resuspended in 95ul of LiCl buffer. The washed M-270 beads were added to 40ul of purified biotinylated cDNA/RNA hybrids. Binding was carried out for 15min at 37C, then beads were purified using a magnetic bar and washed with TE wash buffer (10mM Tris-HCl (pH7.5), 0.1% Tween20, 1mM EDTA(pH8.0)) three times.

Captured cDNA was released from the beads by heat shock and RNaseI treatment. Beads were resuspended in 35ul of release buffer (1x RNaseONE buffer, 0.01% Tween20), incubated at 95C for 5min and chilled on ice immediately. The supernatant containing cDNA was transferred to a new tube. The beads were washed with 30ul of release buffer, and the supernatant was pooled together with the first elution. Then the sample was treated with RNase (0.1ul of 60U/ul RNaseH (TaKaRa) and 2ul of 10U/ul RNaseI for 30min at 37C) to remove RNA completely. Then the Cap-Trapped cDNA was purified with Agencourt AMPure XP (Beckman coulter) according to the manufacture’s protocol. The cDNA quantity was determined with the Quant-iT OliGreen ssDNA Assay kit (ThermoFisher Scientific).

### 5. Linker Ligation

5’/3’ linkers was ligated to the both end of Cap-trapped cDNA.

#### 5.1 How to make a linker

Dissolve the oligonucleotides of 5’ linker to 1mM in TE buffer. For the annealing reaction, GN5 linker reaction solution (4ul of 5’ linker up GN5 (5’- GTGGTAUCAACGCAGAGUACGNNNNN -P-3’: 1mM), 4ul of 5’ linker down (5’-P-GTACTCTGCGTTGATACCAC-P-3’: 1mM), 4 ul of 1M NaCl and 28 ul of water) and N6 linker reaction solution (1ul of 5’ linker up N6 (5’-GTGGTAUCAACGCAGAGUACNNNNNN -P-3’: 1mM), 1ul of 5’ linker down (1mM), 1 ul of 1M NaCl and 7 ul of water) were incubated the following conditions: 95°C, 5 min gradient 0.1°C/sec, 83°C, 5 min, gradient 0.1°C/sec, 71°C 5 min, gradient 0.1°C/sec, 59°C 5 min, gradient 0.1°C/sec, 59°C 5 min, gradient 0.1°C/sec, 47°C 5 min, gradient 0.1°C/sec, 35°C 5 min, gradient 0.1°C/sec, 23°C 5 min, gradient 0.1°C/sec and 11°C Hold. The annealed GN5 linker solution(40ul) and N6 linker solution(10ul) were mixed (5’CTR-Seq linker (100uM)). The 5’CTR-Seq linker (100uM) was diluted to 10uM with 0.1M NaCl (in TE).

For the 3’ CTR-Seq linker, 1ul of 3’ CTR-Seq up (5’-AAAAABBBBBBBBGCAUCGCUGTCTCUTAUACACAUCUCCGAGCCCACGAGAC -P-3’) and 1ul of 3’ CTR-Seq down (5’-GTCTCGTGGGCTCGGAGATGTGTATAAGAGACAGCGATGC -3’), 1 ul of 1M NaCl and 7 ul of wather. Then incubate the mixed solution as same condition as the 5’ linker. After annealing step, the UMI part (BBBBBBBB) was filled with Phusion High-Fidelity DNA polymerase (NEB) and dVTPs(dATP/dGTP/dCTP) instead of dNTPs. After filling reaction, the 3’ linker solution was purified with AMPure XP. Then adjust the concentration to 10uM with 0.1M NaCl in TE buffer.

#### 5.2 5’ SSLL

The cDNA solution was dried up using a SpeedVac (80C for 35min). The pellet was dissolved in 4ul of water. After incubation of cDNA solution at 95C for 5min and chilled on ice for 2min, 1ul of 5’ CTR-Seq linker (10uM), which was incubated at 55C for 5min and chilled on ice, was added. Then 10ul of Mighty Mix (TaKaRa) was added, mixed gently and incubated at 30C for 4h. The sample after ligation was purified with AMPure XP.

#### 5.3 USER

To shorten the long polyT stretch of RT primer, the U residues in the RT primer were digested with USER enzyme (NEB). We added 2ul of USER enzyme (2U/ul), 5ul of 10x CutSmart buffer (NEB) and 3ul of water to 40ul of 5’ linker ligated cDNA. We incubated the reaction solution at 37C for 30min and chilled on ice.

Then the dT stretch at 5’ end of cDNA became 5nt. The cDNA was purified with AMPure XP beads.

#### 5.4 3’ SSLL

The cDNA solution was dried up using a SpeedVac (80C for 35min). The pellet was dissolved in 4ul of water. After incubation of cDNA solution at 95C for 5min and chilled on ice for 2min, 1ul of 3’ CTR-Seq linker (10uM), which was incubated at 65C for 5min and chilled on ice, was added. Then 10ul of Mighty Mix was added, mixed gently and incubated at 16C for 16h. The sample after ligation was purified with AMPure XP.

### 6. SAP treatment

To digest excessed 3’ linker and dephosphorylate the 3’end of 5’linker down strand, the cDNA was treated with 1ul of SAP (Affymetrics) and 2ul of USER in 1x SAP buffer, incubated at 37C for 30min. After reaction, the cDNA was purified with AMPure XP.

### 7. 2^nd^ strand synthesis

The cDNA solution was concentrated to 5ul using a SpeedVac (80C for 35min). The 2^nd^ strand synthesis was carried out using 5ul of cDNA, 0.5ul of 2^nd^ primer (5’ – TCGTCGGCAGCGTCAGATGTGTATAAGAGACAGNNNNNNNNGTGGTATCAACGCAGAGTAC -3’:100uM), 1.3ul of DMSO, 5.8ul of water and 12.5ul of 2x KAPA HiFi HS mix (NIPPON Genetics). The reaction mix was incubated for the following condition; 95C for 5min, 55C for 5min, 72C for 30min and hold at 4C. After 2^nd^ strand synthesis, the excessed primer were digested with adding 1ul of Exonuclease I (20U/ul, NEB) and incubation at 37C for 30min. Then the sample solution was purified with AMPure XP twice. The volume of used AMPure XP beads was 46.8ul at 1^st^ purification and 40ul at 2^nd^ purification. The sample was dried up with SpeedVac (37C for 75min). The pellet was dissolved in 7ul of water.

### 8. quantification/qualification

The ds cDNA was quantified using Quant-iT PicoGreen Assay kit (Thermofisher Scientific), according to the manufacturer’s instructions. For quantification, we used 1ul of ds-cDNA. And we analyzed the length distribution with Agilent High Sensitivity DNA kit (Agilent).

### 9. cDNA amplification

The double stranded cDNAs were amplified using Illumina adapter-specific primers and LongAmp Taq DNA polymerase (NEB). After 16 cycles of PCR (8 minutes for elongation time), amplified cDNAs were purified with equal volume of AMPure XP beads (Beckmann Coulter).

### 10. NanoPore Sequencing

Purified cDNAs were subjected to Nanopore sequencing library following to manufacturer’s 1D ligation sequencing protocol (version NBE_9006_v103_revO_21Dec2016). Nanopore libraries were sequenced by MinION Mk1b with R9.4 flowcell. Sequence data was generated by MinKNOW 1.7.14

### 11. NanoPore Basecalling

In order to generate fastq files from FAST5 files, Basecalling was processed by “Albacore v2.1.0” basecaller software which was provided from Oxford NanoPore Technologies.

### 12. Trimming adapter sequence from fastq file

To preparing clean reads from fastq files, trimming was processed by “Porechop v0.2.3”.

### 13. Method for aligning RIKEN MinION cDNA reads to the human genome

* Software versions: LAST 941, Python 2

First, an index (named “hdb”) of the genome and linkers was prepared:

> lastdb -P0 -uNEAR -R01 hdb hg38.analysisSet.fa linkers.fa

Then, the rates of insertion, deletion, and substitution between reads and genome were estimated:

> last-train -P0 --matsym hdb BC01_A549_OligoDT.fa > f6nano.mat

This was done for BC01 and BC02, with and without --matsym. The results were similar, and the result of the above command was used in the next steps.

The reads were aligned to the linkers:

> lastdb -c -uNEAR linkerdb linkers.fa
>
> echo “N 0 0 0 0” | cat f6nano.mat – |
>
> lastal -P0 -p- linkerdb reads.fa | last-split -m1 > linkers-reads.maf

(This adds a row of zero scores for N to the score matrix, which is appropriate for the UMI with Ns. The other UMI with Vs/Bs is scored appropriately by default.)

Then the reads were oriented in the RNA forward-strand direction:

> analyze-linkers.py reads.fa linkers-reads.maf > reads-fwd.fa 2> linkers-reads.txt

The .txt files have some statistics on linker analysis failures.

Finally, the reads were aligned to the genome:

> parallel-fasta -k “lastal -p f6nano.mat -d90 -m50 -D10 hdb | last-split -g hdb -m1” < reads-fwd.fa > reads.maf

And alternative alignment formats were prepared:

> maf-convert -j1e6 psl reads.maf | grep -v linker > reads.psl
>
> pslToBed reads.psl reads.bed

\## Warnings

* The results include low-confidence alignments. In the maf files, each alignment has a “mismap” probability, which is the estimated probability that it’s aligned to the wrong place.
* There are probably some incorrect alignments to processed pseudogenes. It’s hard to avoid these completely. (There may also be correct alignments to processed pseudogenes.)
* There may be an artifactual tendency for first exons to begin just after AG, and last exons to end just before GT. This is because the spliced alignment method does not treat linkers differently.

**Figure S1:**
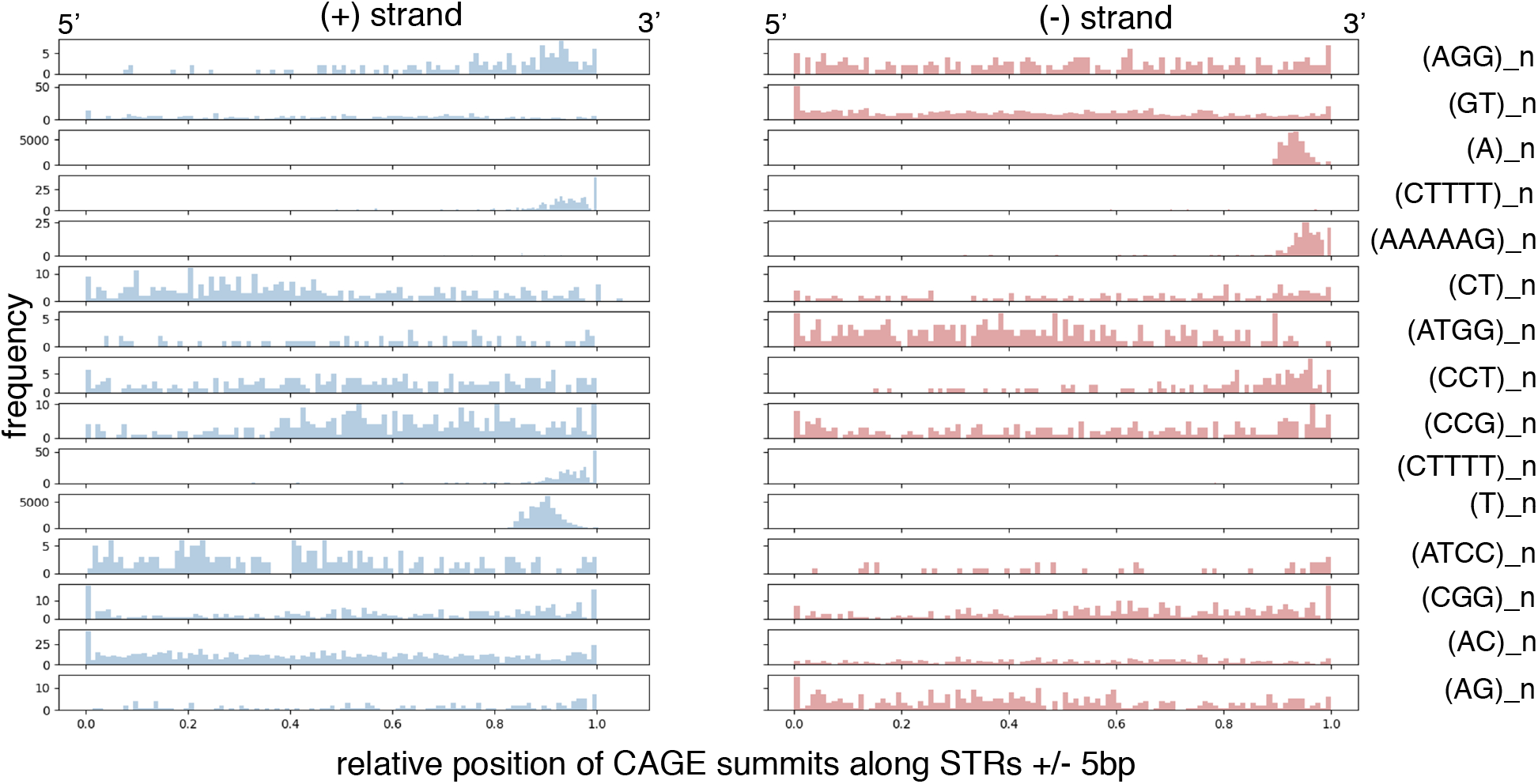
Distribution of CAGE peak summits along STR. The relative position (x-axis) was computed on a window corresponding to STR length ± 5bp. y-axis, frequency of CAGE peaks. Only STR classes with > 200 CAGE peaks on (+) strand and > 200 CAGE peaks on (-) strands are shown.

**Figure S2:**
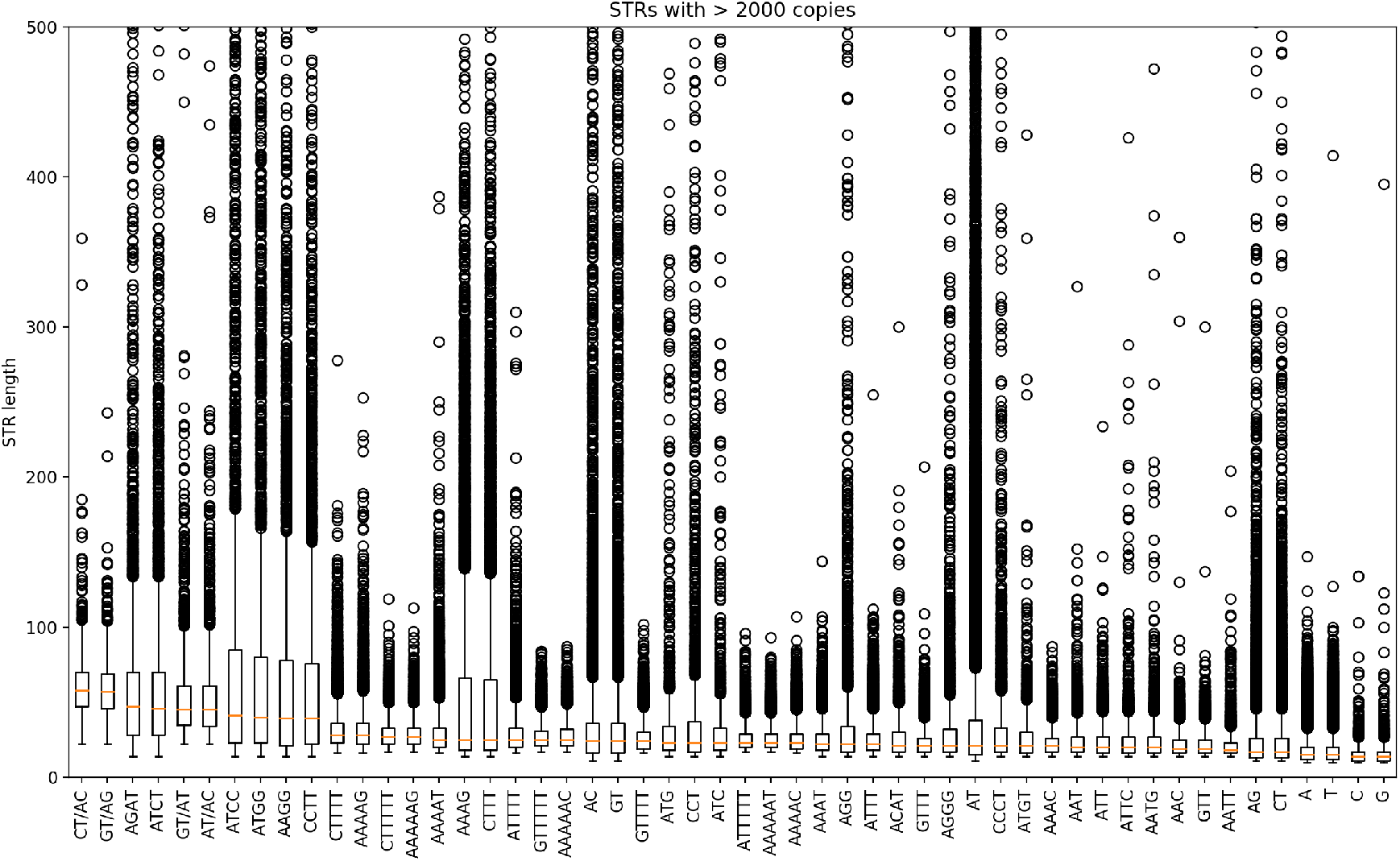
STR length distribution in different classes. STR classes are sorted by median length. Only STR classes with > 2,000 elements are shown.

**Figure S3:**
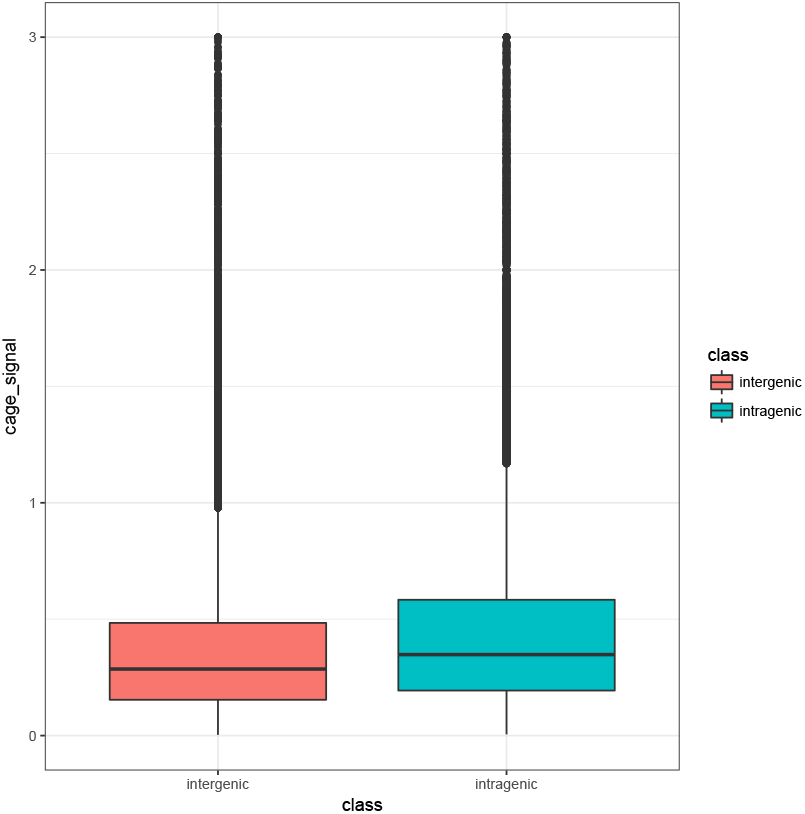
CAGE signal at inter- and intragenic STRs. FANTOM CAT annotation [1] was used to define inter- and intragenic STRs. See Methods section.

**Figure S4:**
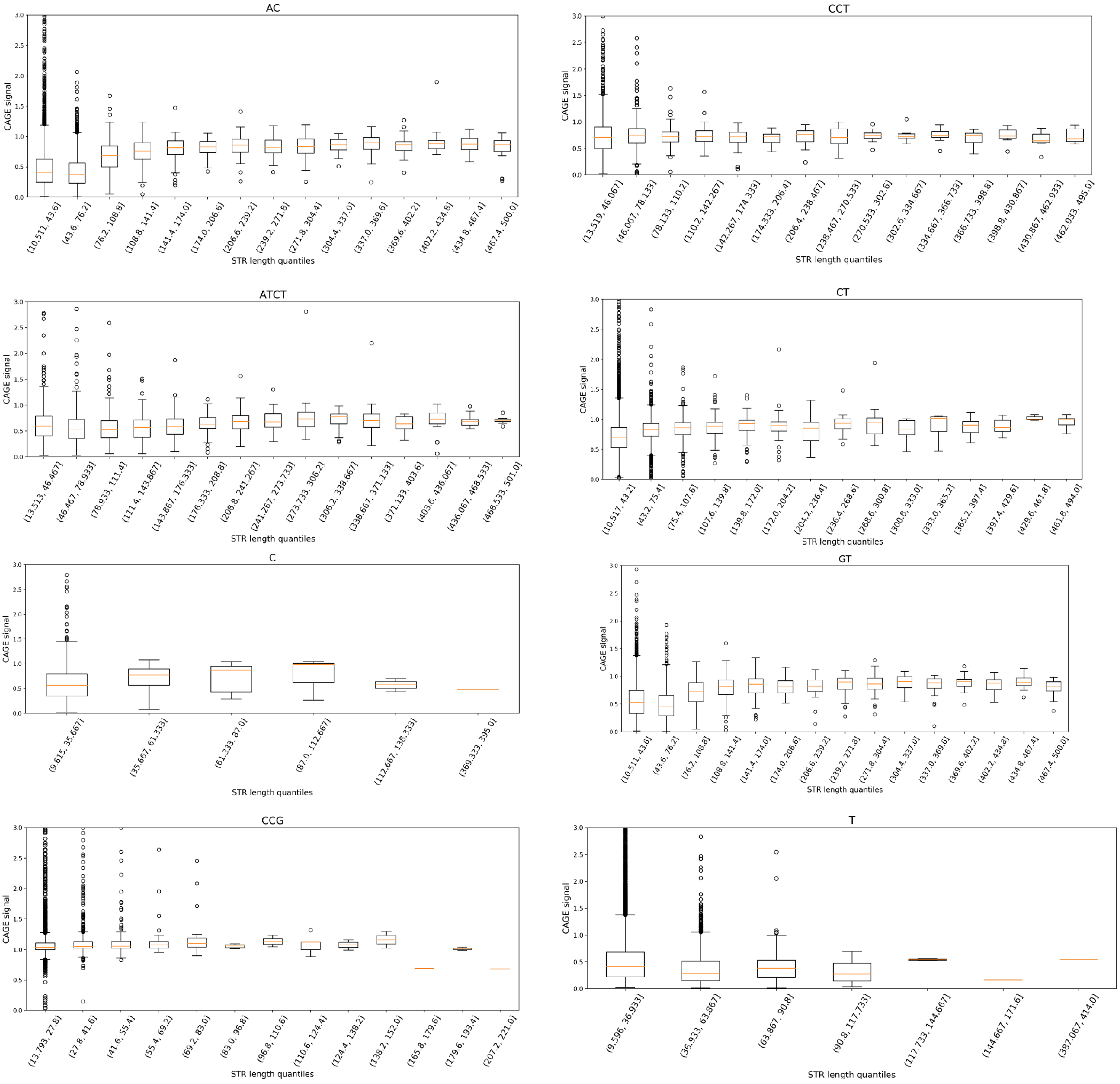
CAGE signal in different STR classes according to STR length. Quantiles were defined using the Pandas quantile-based discretization *qcut* function. x-axis: quantiles; y-axis: CAGE signal

**Figure S5:**
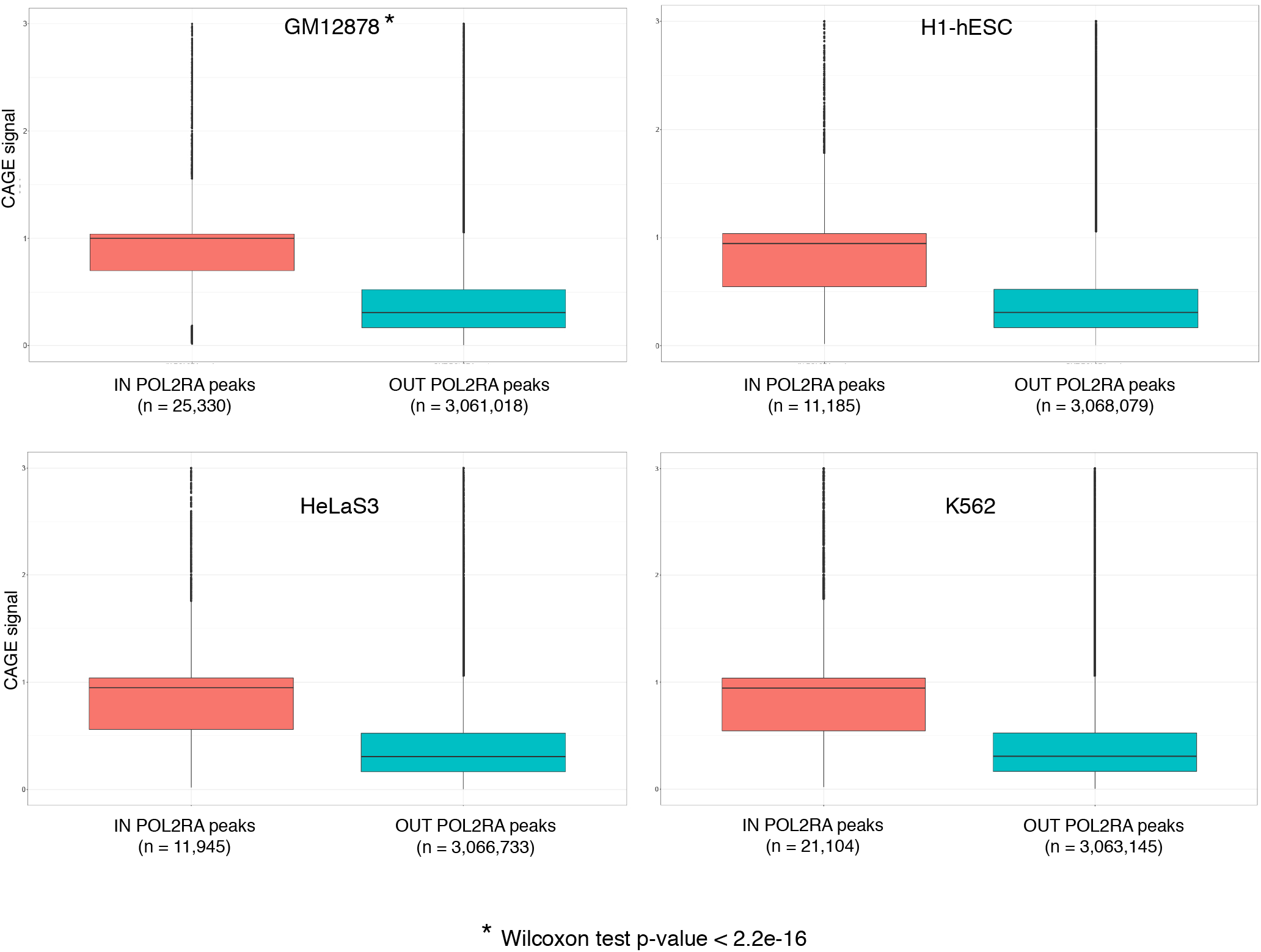
CAGE signal at STRs located within RNAP-II peaks. The coordinates of STRs were intersected with that of RNAP-II ChIP-seq narrow peaks from ENCODE. The CAGE signal associated with STRs located (red) or not (blue) in RNAP-II binding sites were compared. Wilcoxon test was performed in all four cell types tested with a p-value < 2.2e-16.

**Figure S6:**
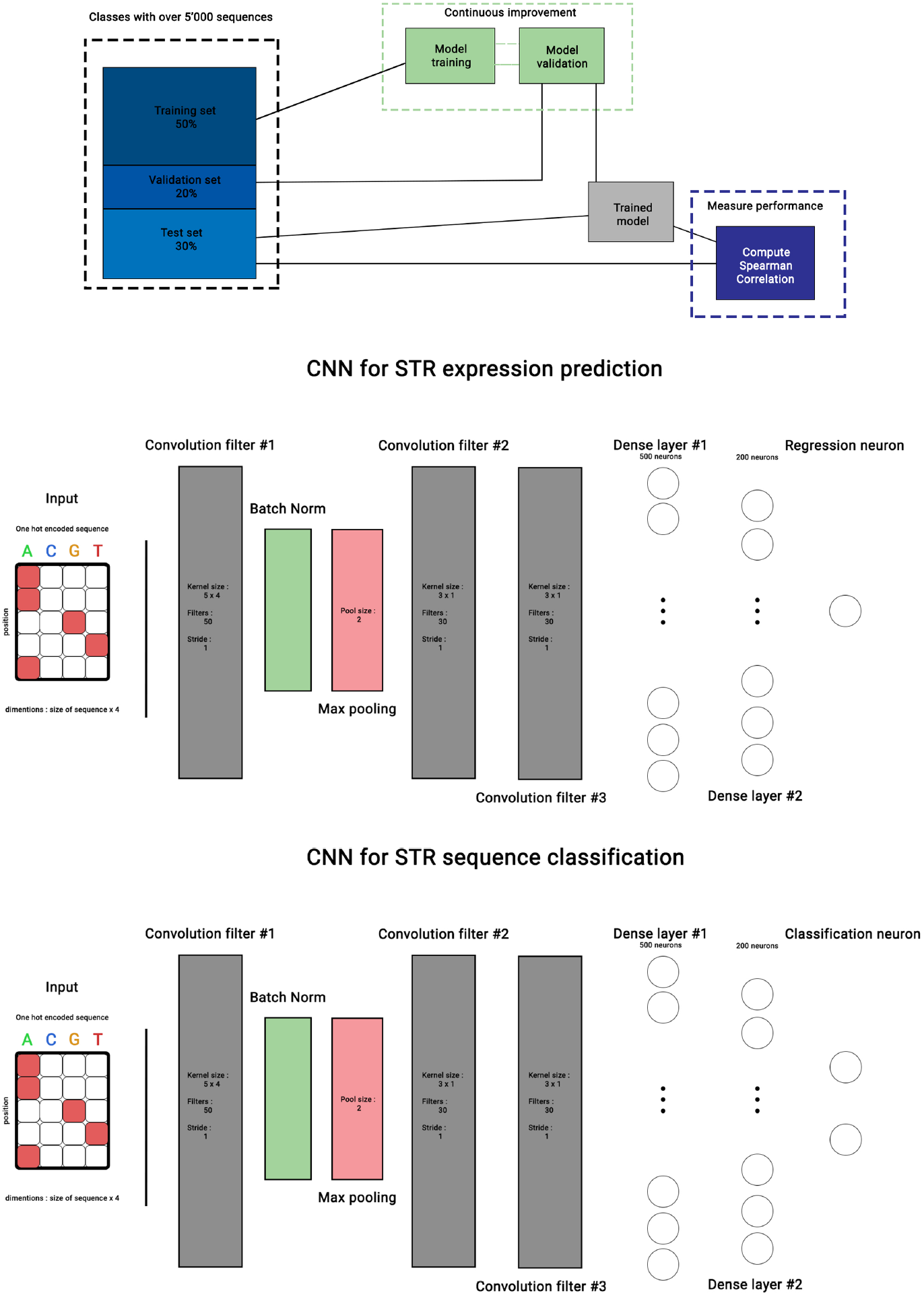
Definition of testing/training sets (top) and model architectures used for classification (bottom) and regression (middle) tasks. The input sequence corresponds to ± 50bp around STR 3’end. Each layer is complemented with a RELU activation function, and dropout is implemented after the first dense layer. Source code is available at https://gite.lirmm.fr/ibc/deepSTR.

**Figure S7:**
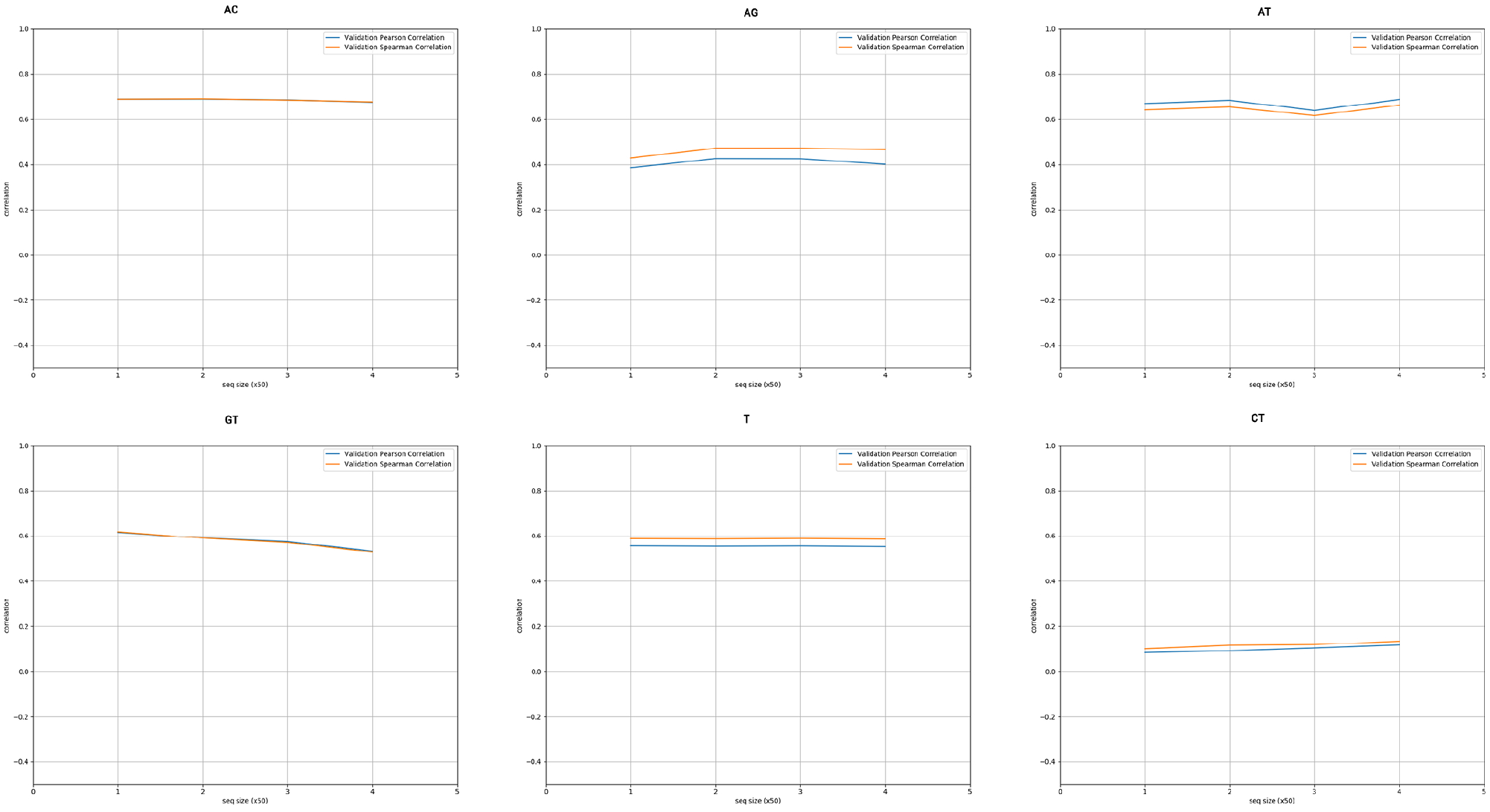
Impact of the length of the sequences used as input of the CNN models. Spearman (orange) and Pearson (blue) correlations (y-axis)were computed between the predicted and the observed CAGE signal. Different sequence size were tested as input (50bp, 100bp, 150bp and 200bp). The size is indicated as multiples of 50bp on the x-axis. Only 6 representative STR classes are shown.

**Figure S8:**
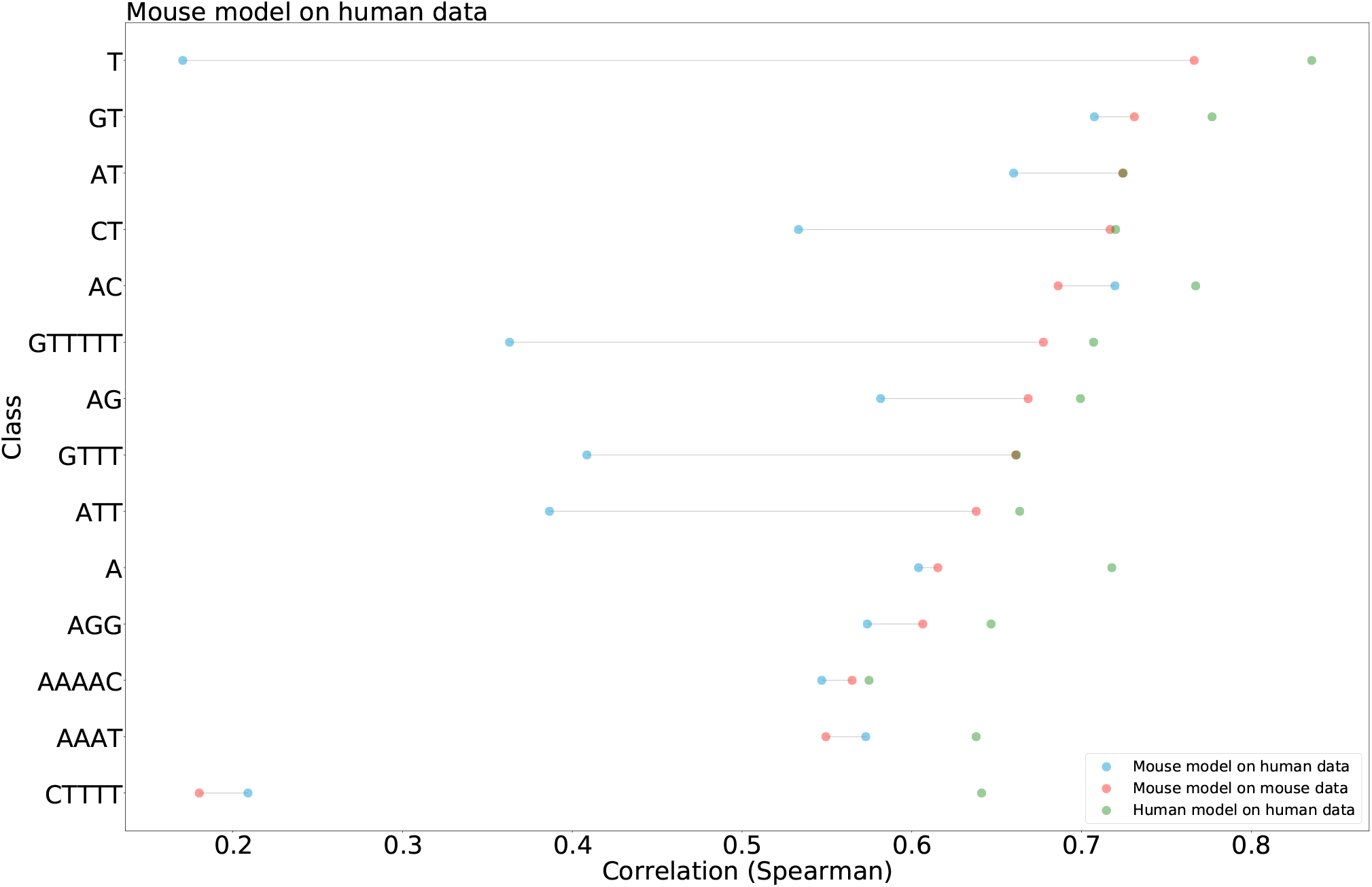
Testing the accuracy of CNN models built in mouse and tested in human for different STR classes. Performance of the model is assessed by computing the Spearman correlation between CAGE signal observed in human and that predicted by a model learned in mouse (blue dots), CAGE signal observed in mouse and that predicted by a model learned in mouse (red dots) and CAGE signal observed in human and that predicted by a model learned in human (green dots). Remember that the mouse models are overall less accurate than human models (Figure 6C). The mouse model is not able to predict transcription at mouse or human (*CTTTT*)_*n*_ (*ρ* < 0.2). Likewise, this model hardly predicts transcription at human (*T*)_*n*_ (*ρ* < 0.2). For other classes, the Spearman correlation between the signal predicted by the mouse model and the observed human signal was > 0.3, confirming that several features are conserved between human and mouse.

**Figure S9:**
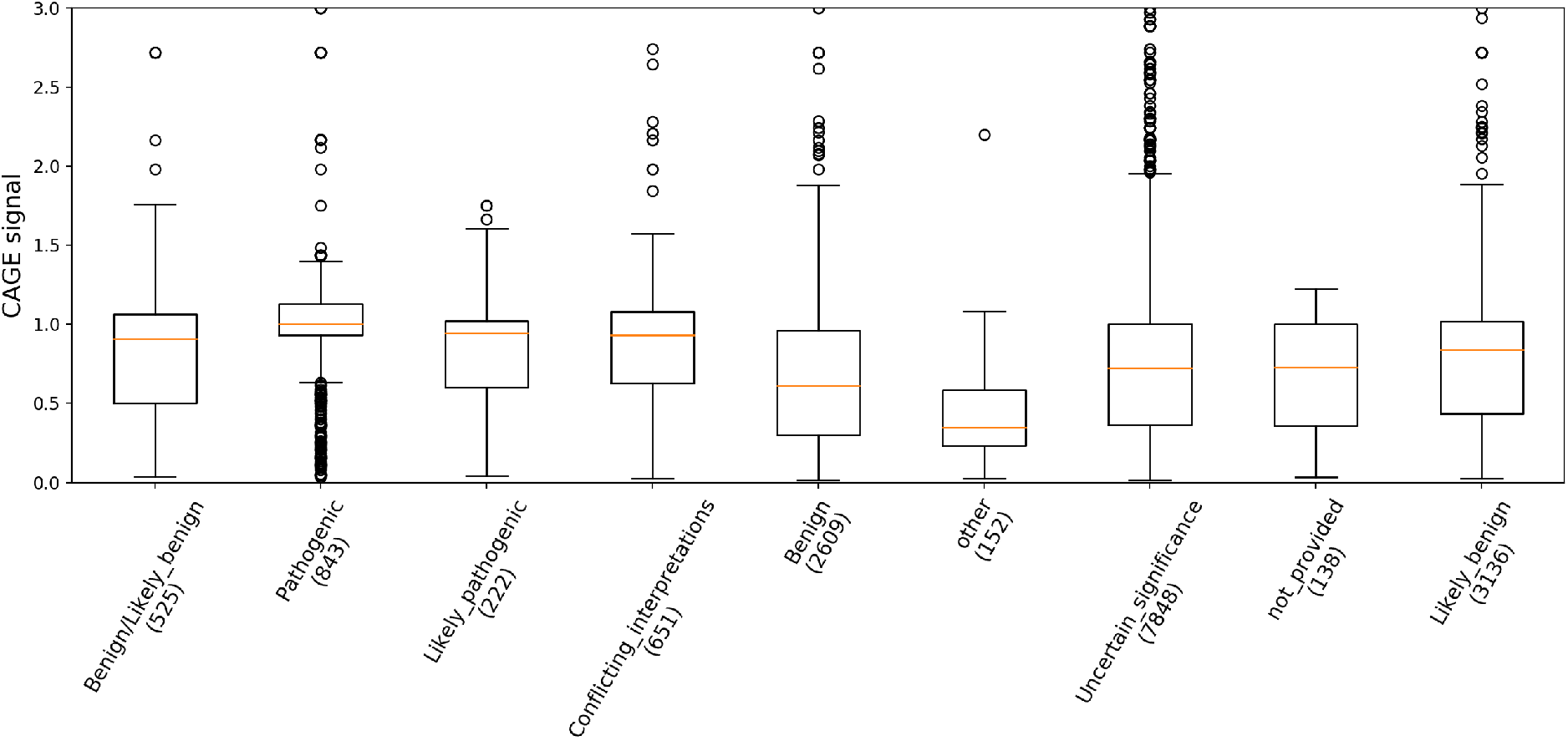
CAGE signal (y-axis) at STRs associated with ClinVar variants ordered according to their clinical significance (x-axis). The number of variants considered for each ClinVar class is indicated in bracket.

**Figure S10:**
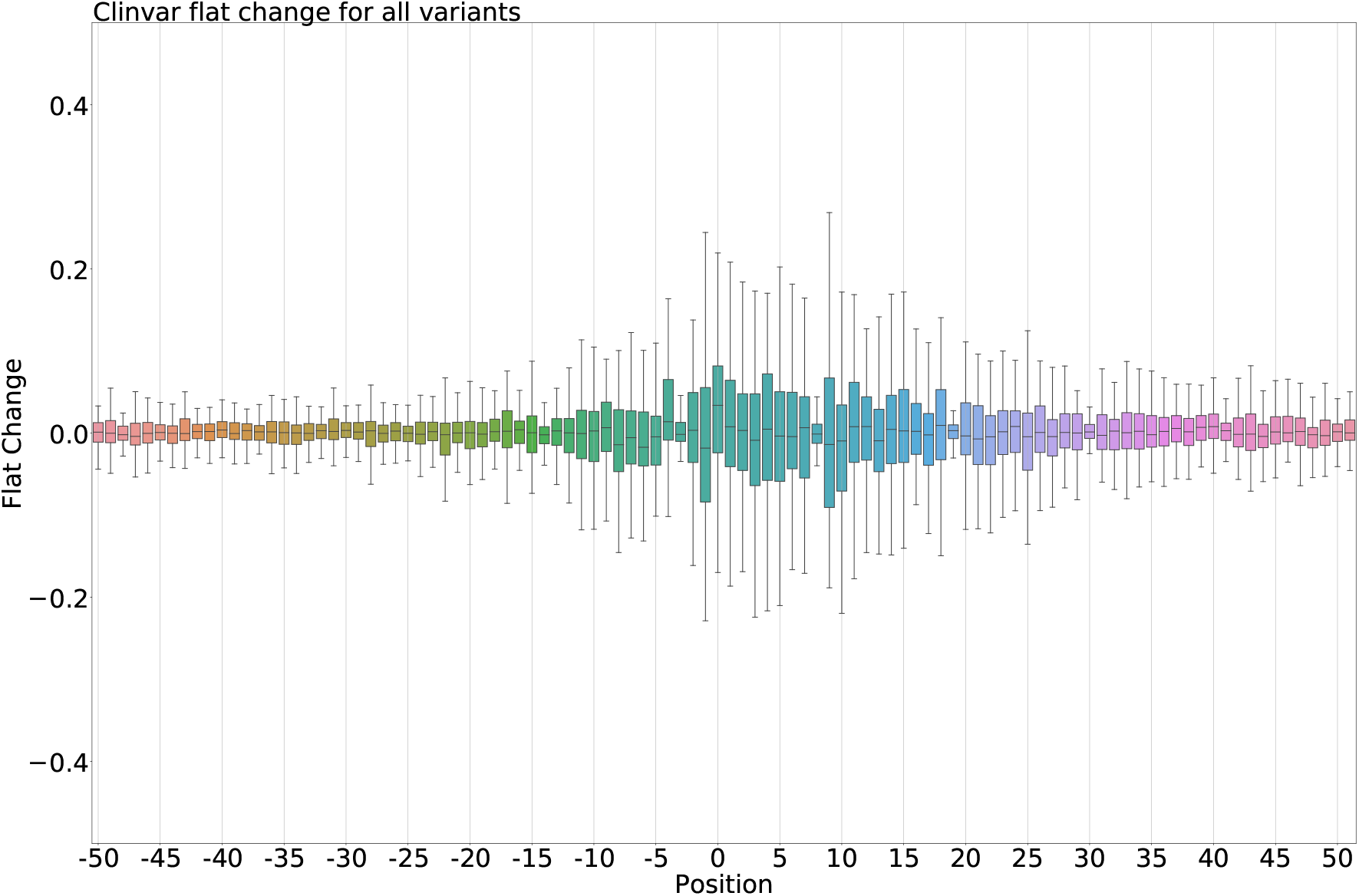
Impact of ClinVar variants on CNN predictions. Predictions are made on the hg19 reference sequence and on a mutated sequence, containing the genetic variants. Note that to keep sequences aligned, only single nucleotide variants are considered. Changes (y-axis) are then measured as the difference between these two predictions (reference – mutated). Values are grouped by the position of the variants relative to the STR 3’ end (position 0 on the x-axis). Note that variations at −3, 8 and 19 have no impact, revealing the potential existence of ‘blind’ positions, where models did not learn features.

**Figure S11:**
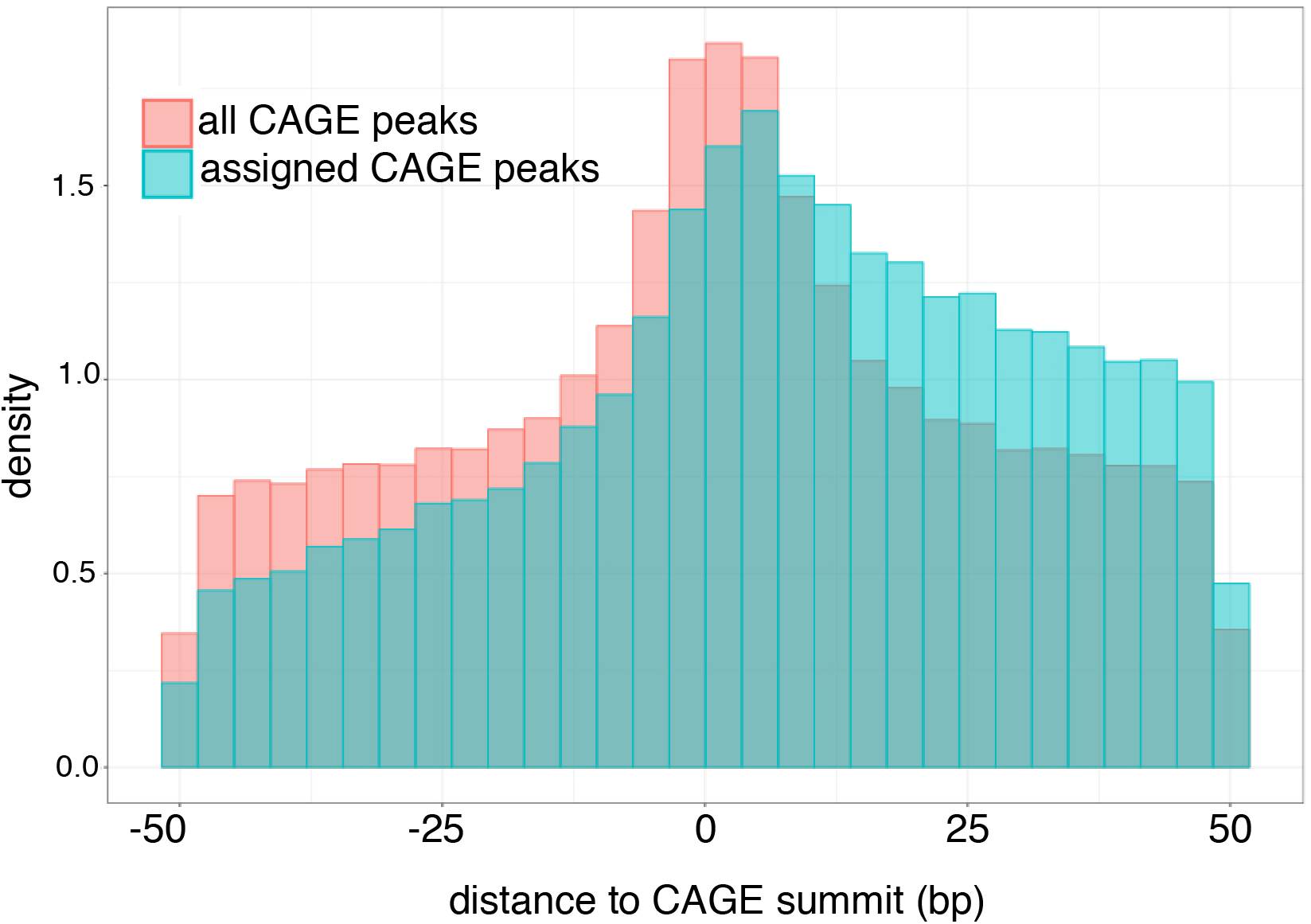
Distribution of ClinVar variants around all CAGE peak summits (red, n = 1,048,124) and CAGE peak summits assigned to genes (blue, n = 130,286).

